# The Alzheimer’s disease risk gene BIN1 modulates neural network activity via the regulation of L-type calcium channel expression in human-induced neurons

**DOI:** 10.1101/2022.01.18.476601

**Authors:** Orthis Saha, Ana Raquel Melo de Farias, Alexandre Pelletier, Dolores Siedlecki-Wullich, Johanna Gadaut, Bruna Soares Landeira, Arnaud Carrier, Anaïs-Camille Vreulx, Karine Guyot, Amelie Bonnefond, Philippe Amouyel, Cláudio Marcos Queiroz, Devrim Kilinc, Fabien Delahaye, Jean-Charles Lambert, Marcos R. Costa

## Abstract

**Background:** Bridging Integrator 1 (*BIN1*) is the second most important Alzheimer’s disease (AD) risk gene, but its physiological roles in neurons and its contribution to brain pathology remain largely elusive. In this work, we show that BIN1 plays a critical role in the regulation of calcium homeostasis, electrical activity, and gene expression of glutamatergic neurons.

**Methods:** We generated 3D cerebral organoids and 2D enriched neuronal cell cultures from isogenic *BIN1* wild-type (WT), heterozygous (HET) and homozygous knockout (KO) human-induced pluripotent stem cells (hiPSCs). Using single-cell RNA-sequencing, biochemical assays, immunocytochemistry and multi-electrode array(MEA) electrophysiology, we characterized the molecular and functional consequences of reduced BIN1 expression in different neural cell types.

**Results:** We show that *BIN1* is mainly expressed by oligodendrocytes and glutamatergic neurons of cerebral organoids, like in the human brain. Both *BIN1* HET and KO cerebral organoids show specific transcriptional alterations, mainly associated with ion transport and synapses in glutamatergic neurons. We then demonstrate that *BIN1* cell-autonomously regulates gene expression in glutamatergic neurons by using a novel protocol to generate pure culture of human-derived induced neurons (hiNs). Using this system, we also show that BIN1 plays a key role in the regulation of neuronal calcium transients and electrical activity via its interaction with the L-type voltage-gated calcium channel Cav_1.2_. *BIN1* KO hiNs show reduced activity-dependent internalization and higher Cav_1.2_ expression compared to WT hiNs. Pharmacological treatment with clinically relevant doses of nifedipine, a calcium channel blocker, partly rescues neuronal electrical and gene expression alterations in *BIN1* KO glutamatergic neurons. Further, we show that transcriptional alterations in *BIN1* KO hiNs affecting biological processes related to calcium homeostasis are also present in glutamatergic neurons of the human brain at late stages of AD pathology.

**Conclusions:** Together, our findings suggest that BIN1-dependent alterations in neuronal properties could contribute to AD pathophysiology and that treatment with low doses of clinically approved calcium blockers should be considered as an option to dampen disease onset and progression.

## Introduction

The Bridging Integrator 1 (*BIN1*) is the second most associated genetic determinant with the risk of late-onset Alzheimer’s disease (LOAD), after the Apolipoprotein E (*APOE*) gene [1–4]. In the adult human brain, *BIN1* is mainly expressed by oligodendrocytes, microglial cells and glutamatergic neurons [5–7] and its expression is reduced in AD patients compared to healthy individuals [7,8]. The consequences of this reduced BIN1 expression to neuronal and glial cells, as well as the mechanisms by which it contributes to AD pathogenesis remain poorly understood.

BIN1 has disputably been associated with amyloidopathy and tauopathy, two pathological hallmarks of AD [9]. Reduced BIN1 expression results in a higher amyloid precursor protein (APP) processing towards the production of amyloid-beta (Aβ) peptides in Neuroblastoma Neuro2a cells [10,11]. However, we previously showed that *BIN1* knockout (KO) does not increase the concentrations of Aβ peptides in hiPSC-derived neurons (human-induced neurons or hiNs) despite impairing endocytic trafficking [12]. Likewise, reduced Bin1 expression in the mouse brain does not affect the production of endogenous Aβ peptides [13]. Concerning Tau pathology, decreased expression of the BIN1 ortholog Amph suppress Tau-mediated neurotoxicity in Drosophila Melanogaster [14]. In contrast, reduced BIN1 expression results in higher Tau aggregation and propagation in primary rat hippocampal neurons [15]. In humans, higher concentrations of phosphorylated Tau are observed in the cerebrospinal fluid of patients with AD and are significantly correlated with genetic variants within the *BIN1* locus [16].

More recently, BIN1 has also been associated with the regulation of synaptic transmission and neuronal electrical activity in animal models. Conditional deletion of *Bin1* in neurons of the adult mice hippocampus leads to altered frequency of mini excitatory post-synaptic currents (mEPSC), likely due to an impaired presynaptic release probability and slower depletion of neurotransmitters [6]. Knockdown of *BIN1* in embryonic rat primary cortical neurons also affects the glutamate AMPA receptor trafficking in the post-synaptic compartment, leading to alterations in the amplitude of mEPSC [17]. Lastly, overexpression of a BIN1-mKate2 fusion protein increases the frequency of spontaneous excitatory postsynaptic currents (sEPSCs) in embryonic rat hippocampal cultures, seemingly by affecting the localization of L-type voltage gated calcium channels (LVGCC) in the membrane through a Tau-dependent interaction [18].

Despite these advances, no consensus has been reached on the roles of BIN1 in AD pathogenesis and even its physiological functions in human brain cells remain mostly unknown. In this work, we tackled this important question by generating and characterizing human neural cells derived from isogenic *BIN1* wild type (WT), heterozygous (HET) and homozygous knockout (KO) hiPSC lines. We first characterized the transcriptional profile of human neural cells grown in three-dimensional cerebral organoids for more than 6 months and show that reduced BIN1 expression affects mainly glutamatergic neurons. Next, we generated pure *BIN1* WT and KO hiNs cultures and show that BIN1 cell-autonomously regulates electrical activity and gene expression of glutamatergic neurons via the interaction with the regulation of the LVGCC Cav_1.2_. Pharmacological blockage of this channel with nifedipine partly rescues electrical and gene expression alterations in *BIN1* KO hiNs. Our findings suggest that BIN1 is a key regulator of calcium homeostasis in glutamatergic neurons and that repurposing the use of clinically approved calcium channel blockers could be a promising strategy to treat AD.

## Results

### BIN1 HET and KO cerebral organoids show transcriptional alterations associated with neuronal functional properties

Cerebral organoids (COs) faithfully recapitulate fundamental aspects of the three-dimensional organization of the human brain, including the molecular specification of different neural cell types/subtypes and the generation of complex electrical activity patterns [19,20]. To investigate the potential role of BIN1 in human neural cells, we generated and characterized COs using isogenic *BIN1* wild type (WT), heterozygous (HET) and KO hiPSCs (Fig. 1A). After 6.5 months of culture, COs were composed of all the major neural cell types identified by the expression of MAP2, GFAP and NESTIN and we did not observe any gross difference in size or morphology of COs among the three genotypes (Sup. Fig. 1A). Western blot analyses confirmed the reduction and absence of BIN1 protein in BIN1 HET and KO COs, respectively (Sup. Fig. 1B). We have then employed single-nucleus RNA sequencing (snRNA-seq) to further characterize individual cell types/subtypes and investigate possible gene expression alterations associated with reduced BIN1 expression. COs (n=4 from each genotype) were divided into two halves that were independently processed for western blotting or snRNA-seq. We observed similar expression of general neuronal and glial proteins in these COs (Sup. Fig. 1C), suggesting a low degree of heterogeneity in these samples. Nevertheless, to further reduce potential batch effects, we pooled COs into a single multiplexed library using Cell Hashing (Stoeckius et al., 2018). After sequencing, quality control and demultiplexing, we recovered 4398 singlets that could be grouped into 7 major cell clusters based on the expression of cell type markers *SLC1A3 (GLAST), GFAP* and *TNC* (astrocyte); *SNAP25, DCX, MAPT* (pan-neuronal); *SLC17A7* and *SLC17A6* (glutamatergic neurons); *DLX1, GAD1* and *GAD2* (GABAergic neurons); *HES6, CCND2* and *CDK6* (NPCs); *ITGA8* (choroid plexus); and *CLIC6* (pigmented epithelium) (Fig. 1B-C). *BIN1* expression in COs was mainly detected in glutamatergic neurons and oligodendrocytes (Fig. 1C), similar to the profile described for the human brain [7] - except for brain microglial cells that express *BIN1* but are not present in COs. We also observed a reduction and enlargement, respectively, in the proportions of glutamatergic neurons and astrocytes in *BIN1* KO compared to WT (Fig. 1D; ****p<0.0001; Chi-square test).

**Figure 1:**
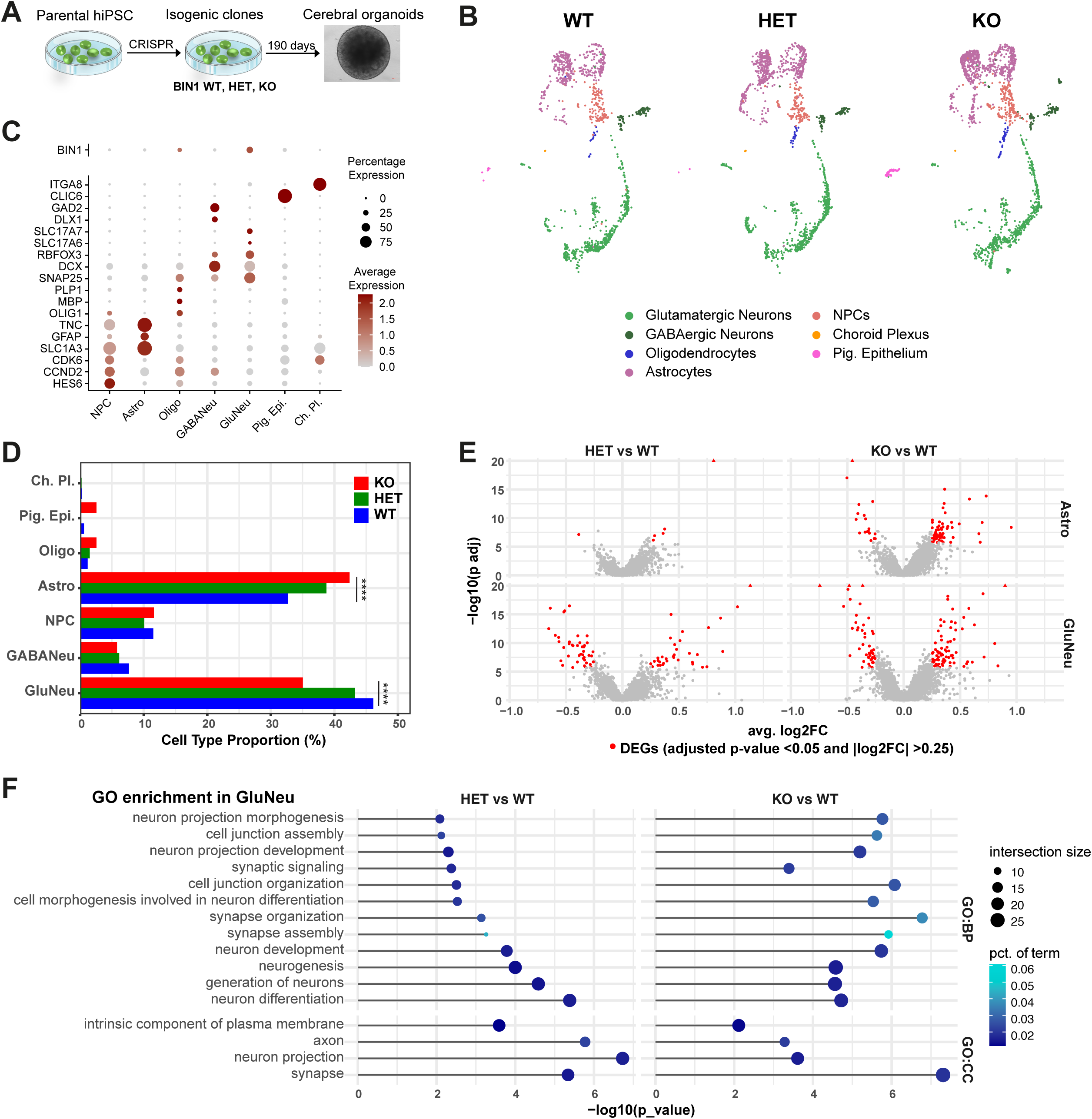
Similar transcriptional changes in glutamatergic neurons from 6.5-month-old BIN1 HET and KO COs. (A) Scheme of the experimental design. (B) UMAP representation of the different cell subtypes in COs identified using snRNA-seq. (C) Dot plot representing the expression for *BIN1* and key markers used to annotate cell subtypes. (D) Proportion of cell subpopulations in the three genotypes (****p<0.0001; Chi-squared test). (E) Volcano plots representing DEGs comparing HET vs WT or KO vs WT in astrocytes and glutamatergic neurons. DEGs with adjusted p-value <0.05 and |log2FC| >0.25 are shown in red. (H) Functional enrichment analysis of DEGs commonly identified in HET and KO glutamatergic neurons. GO: gene ontology; BP: biological processes; CC: cellular components; MF: molecular function.

To identify possible differentially expressed genes (DEGs) in *BIN1* KO or HET compared to WT cells, we performed Wilcoxon test for each major cell type/subtype identified in COs. Consistent with the predominant expression of *BIN1* in glutamatergic neurons (Fig. 1D), we identified a high number of DEGs in this cell type both in *BIN1* HET (76 genes) and KO (124 genes) compared to WT genotype (Fig. 1E; Sup. Fig. 1D). In astrocytes, we also detected 75 DEGs in *BIN1* KO, but only 6 DEGs in *BIN1* HET compared to WT (Fig. 1E; Sup. Fig. 1D). For all other cell types, we observed a maximum of 1-4 DEGs in the comparison between KO vs WT or HET vs WT (Sup. Table 1). These observations suggest that both BIN1 null (KO) and partial deletion (HET) affect similar biological process in glutamatergic neurons in a dose-dependent manner. Accordingly, similar GO terms were enriched for DEGs identified in *BIN1* KO or HET glutamatergic neurons, including several terms associated with synaptic transmission (Fig. 1F). For *BIN1* KO glutamatergic neurons, we also identified GO terms associated with ion channel complex and calcium ion binding (Sup. Fig. 1E; Sup. Table 2), further suggesting that BIN1 depletion leads to specific transcriptional changes associated with functional properties of this neuronal subtype.

### Altered expression of activity-related genes in BIN1 KO and HET COs

Neuronal firing patterns (such as tonic and burst firing) play a key role on the transcriptional regulation of a particular set of genes designated activity-related genes (ARGs) [21]. While neurons stimulated with brief patterns of electrical activity transcribe rapid primary response genes (rPRGs) or early response genes (ERGs), those stimulated with sustained patterns of electrical activity express delayed primary response genes (dPRGs), secondary response genes (SRGs) or late response genes (LRGs) (Fig. 2A)[22,23]. Using Cell-ID [24], we quantified the enrichment for ARGs signatures (Sup. Table 3) in COs at single-cell resolution as an indirect read-out of neuronal electrical activity patterns in this model. We first confirmed that ARG signatures were predominantly enriched in neurons (Fig. 2B). Then, we quantified the proportion of glutamatergic or GABAergic neurons significantly enriched for such specific response gene signatures (p_adj_<0.05; hypergeometric test). We observed a significantly higher proportion of glutamatergic neurons enriched for dPRGs and LRGs both in *BIN1* HET and KO, as well as SRGs in *BIN1* KO compared to WT glutamatergic neurons, whereas the proportion of glutamatergic neurons enriched for rPRGs and ERGs was reduced in *BIN1* HET and KO (Fig. 2C). In sharp contrast, the only difference observed in GABAergic neurons was a reduction in the proportion of cells enriched for SRGs (Sup. Fig. 2). These results suggest that reduced BIN1 expression in glutamatergic neurons triggers neuronal firing patterns towards sustained activity leading to a higher expression of late-response ARGs.

**Figure 2:**
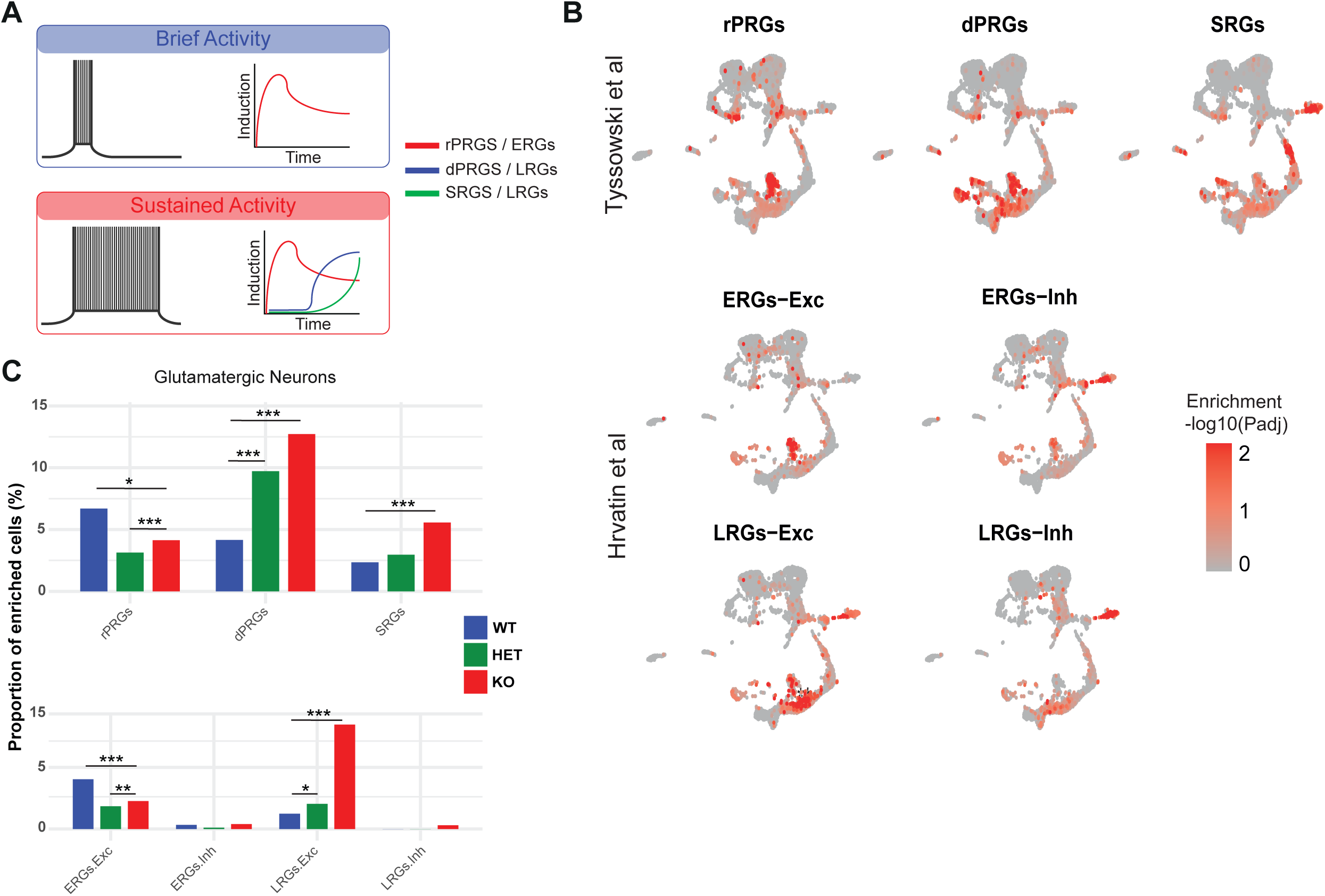
Altered expression of activity-related genes in BIN1 HET and KO glutamatergic neurons. (A) Scheme indicating the different sets of ARGs regulated by brief and sustained patterns of electrical activity [22,23]. rPRGs: rapid primary response genes; dPRGs: delayed primary response genes; SRGs: secondary response genes; ERGs: early response genes; LRGs: late response genes; Exc – glutamatergic neurons; Inh – GABAergic neurons. (B) Feature plots showing the enrichment score of single cells for ARG signatures. Enrichment scores correspond to the –log10(p_adj_) of the Cell-ID-based enrichment test. (C) Proportions of glutamatergic neurons (right) enriched for the different ARG signatures according to genotype (*p<0.05; **p<0.01; ***p<0.001; Chi-squared test).

### Lower numbers of synaptic puncta in BIN1 HET and KO COs compared to WT

Transcriptional alterations in *BIN1* HET and KO glutamatergic neurons are also suggestive of synaptic dysfunction, which is an early hallmark of AD pathology [25]. We thus sought to determine whether BIN1 depletion could affect synaptic connectivity in COs. Using immunohistochemistry to detect the expression of the pre-synaptic protein Synaptohysin-1 (SYP) and post-synaptic protein HOMER1, we were able to quantify the frequency of putative synaptic contacts (% SYP assigned) in COs (see methods). We observed a significant reduction in the % of SYP assigned both in *BIN1* HET and KO compared to WT, mainly due to a reduction in the number of post-synaptic spots expressing HOMER1 (Fig. 3A-D).

**Figure 3:**
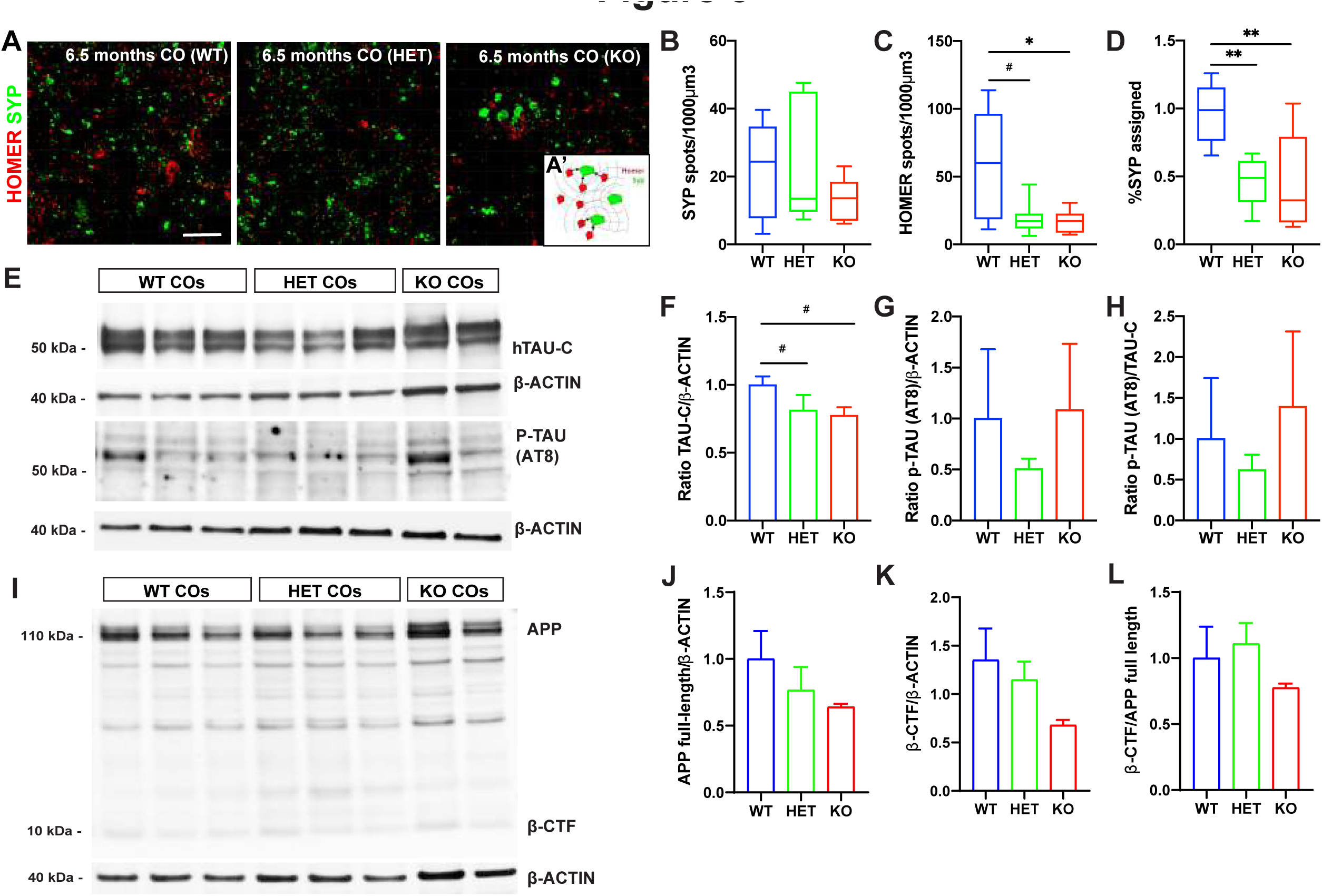
Reduced synaptic numbers in BIN1 HET and KO cerebral organoids. (A) Immunohistochemistry for HOMER1 (red), SYP (green) in 6.5-month-old *BIN1* WT, HET and KO COs. Insert (A’) illustrates the identification of putative synaptic contacts. (B-D) Quantifications of the SYP and HOMER1 spot density, and the percentage of SYP assigned by HOMER1 spots in *BIN1* WT, HET and KO COs (^#^p<0.1; *p<0.05; **p<0.01; Dunn’s multiple comparison test; n=3 COs per genotype). (E) Western blot for total TAU protein C-terminal (TAU-C), phosphorylated (p)-TAU at Ser202, Thr205 (AT8) and β-ACTIN in 6.5-month-old COs. (F-H) Quantification of TAU-C/β-ACTIN, p-TAU/β-ACTIN and p-TAU/TAU-C levels normalized to WT (^#^p<0.1; Mann-Whitney test; n=3 COs per genotype). (I) Western blot for APP and β-ACTIN in 6.5-month-old COs. (J-L) Quantification of APP/β-ACTIN, β-CTF/β-ACTIN and β-CTF /APP levels normalized to WT (n=3 COs per genotype).

Next, we investigated whether APP processing and Tau phosphorylation, which have been previously associated with BIN1 and are known to modulate neuronal electrical activity [10,15,26], could also be altered in *BIN1* HET and KO COs. To this end, we measured the intracellular levels of full-length APP and APP β-CTF (as a readout of amyloidogenic APP processing), total and phosphorylated TAU proteins by western blotting. Besides a trend for reduced TAU expression (Fig. 3F) likely explained by the reduced proportion of neurons, we did not detect any significant differences in the intracellular levels of APP, APP β-CTF, TAU or phospho-TAU (Ser202, Thr205) in *BIN1* HET and KO compared to WT COs (Fig. 3E-L). Altogether, these results suggest that BIN1 depletion could alter neuronal functional properties without significantly affecting APP or Tau metabolism in COs.

### Cell-autonomous role of BIN1 in the regulation of neuronal gene expression

Our results in COs suggest that reduced BIN1 expression affects mainly glutamatergic neurons. However, at least in BIN1 KO COs, we cannot completely rule out an effect of BIN1 deletion in astrocytes that could indirectly impact glutamatergic neurons. Therefore, to unambiguously probe the cell-autonomous effect of *BIN1* deletion on the electrical activity and gene expression of human glutamatergic neurons, we generated *BIN1* WT or KO pure neuronal cultures by direct lineage-reprogramming of human NPCs (hNPCs) using doxycycline-inducible expression of ASCL1 (Fig. 4A; see online methods). After validation of the efficient lineage-reprogramming of hNPCs into highly pure neurons (hereafter ASCL1-hiNs) (Fig. 4B), we added exogenous human cerebral cortex astrocytes to trophically support functional neuronal maturation and synaptic connectivity [27]. Using snRNA-seq after 4 weeks of differentiation we identified 5583 cells (n=5 independent culture batches) clustered into two main glutamatergic neuron (GluNeu-I and II), one GABAergic neuron (GABANeu), one immature/unspecified neuron (UnspNeu), two astrocyte (Astro-I and II) and 1 proliferative NPC groups (Fig. 4C-D). Sample-level differential gene expression analysis using DESeq2 [28], revealed 99 DEGs (|log2FC| >0.25 and FDR <0.05) in *BIN1* KO GluNeu-II compared to WT, but only 2 in GluNeu-I and 1 in immature neurons (Fig. 4E; Sup. Table 4). As observed in COs (Fig. 1H-I), GO term enrichment analysis revealed a significant enrichment for terms associated with synaptic transmission, ion channel activity and calcium signaling pathways (Fig. 4F; Sup. Table 5). The percentage of GluNeu-II enriched for late-response ARGs was slightly greater in *BIN1* KO compared to WT, but without statistical differences (Fig. 4G-H). Exogenously added human astrocytes co-cultured with *BIN1* WT and KO hiNs also showed a low number of DEGs (11 in Astro-I; Sup. Table 4), likely reflecting an astrocyte reaction to primary changes in hiNs in response to *BIN1* deletion.

**Figure 4:**
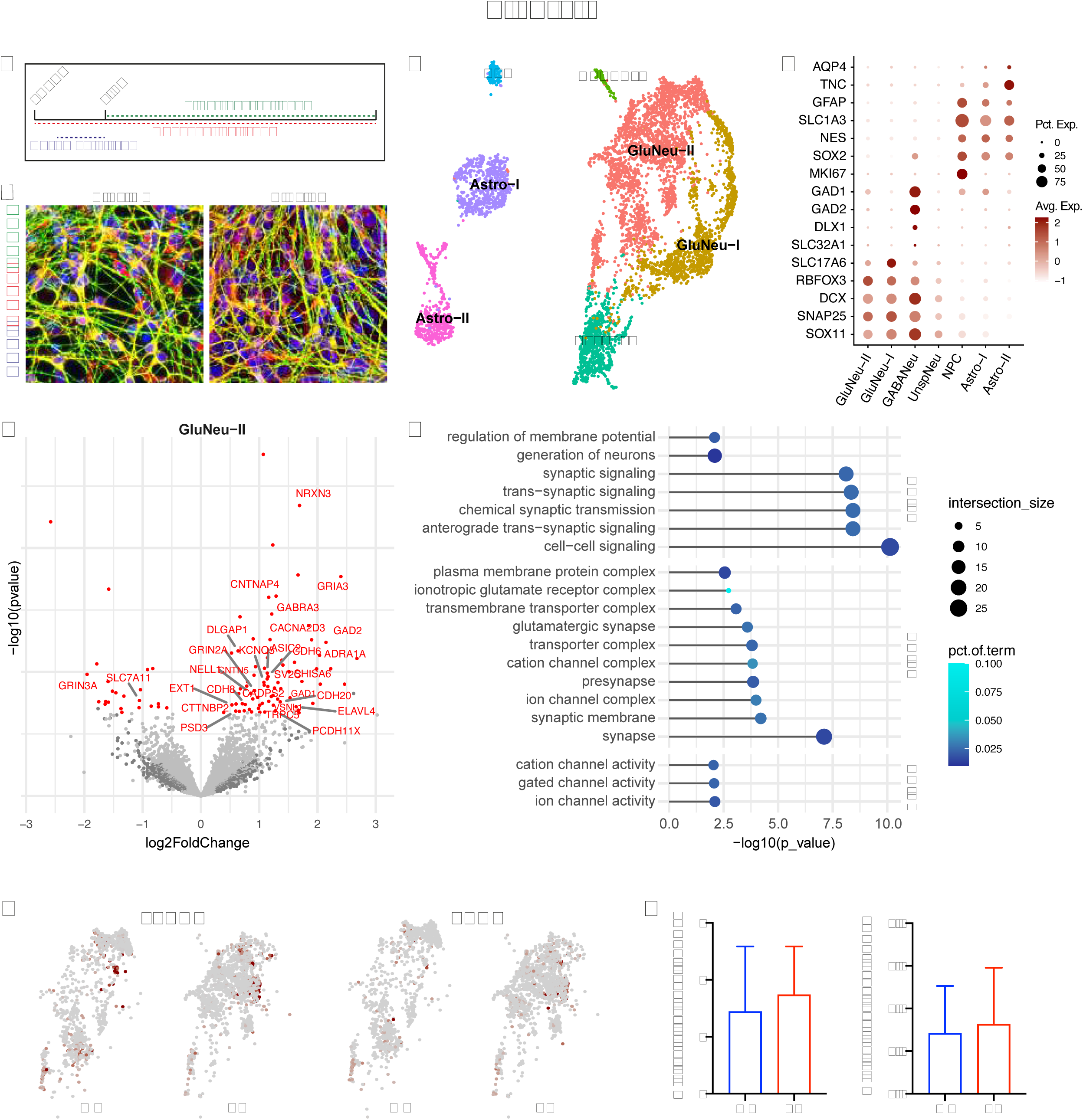
Neuronal-specific BIN1 KO cell-autonomously elicit transcriptional changes. (A) Scheme of the Ascl1-induced hiNs experiments. (B) Images showing BIN1 WT and KO hiNs 7 days after the beginning of doxycycline treatment immunolabeled for neuronal markers MAP2 and TUBB3 and astrocyte marker GFAP and stained with DAPI. (C) UMAP representation of the different cell subtypes identified in ASCL1-hiNs cultures using snRNA-seq. (D) Dot plot representing expression of key markers used to annotate cell subtypes. (E) Volcano plot representing DEGs comparing BIN1 KO vs WT glutamatergic neurons (GluNeu-II). Differential expression analysis was performed using DEseq2 on the sample level gene expression matrix, including the experiment batch as covariates. DEGs with adjusted p-value <0.05 and |log2FC| >0.25 are shown in red. Gene labels are shown for calcium- and synapse-related genes. (F) Functional enrichment analysis of DEGs identified in glutamatergic neurons. Bar plots representing the top 15 the enriched GO terms in each category at adjusted p-value <0.01. (G) UMAP representing GluNeu-II enrichment for delayed Primary Response Genes (dPRGs) and Secondary Responses Genes (SRGs) (Tyssowski et al., 2018). Red point indicate enrichment with adjusted p-value<0.05, hypergeometric test using CellID. (H) Barplot of the percentage of enriched GluNeu-II cells by genotype for each gene sets.

### BIN1 KO leads to alteration in the electrical activity pattern of ASCL1-hiNs

The transcriptional changes observed in our 2D and 3D models could suggest that reduced BIN1 expression is associated with altered electrical properties of glutamatergic neurons. To directly address this possibility, we used multi-electrode arrays (MEA) to record and quantify multi-unit activity (MUA) in ASCL1-hiNs. As previously described in spontaneously differentiated hiPSC-derived neuronal cultures [29], ASCL1-hiNs cells exhibited a diverse range of spontaneous activity patterns, including regular discharges, population bursts and period activity (Fig. 5A). In this respect, we found a conspicuous change in the temporal organization of MUA after *BIN1* deletion (Fig. 5A), mainly characterized by a greater number of spike bursts at 4 weeks (Fig. 5D). These alterations may result from changes at the single cell or the population level (different number of neurons contributing to each electrode, for example). To disentangle these possibilities, we used waveform-based spike sorting to examine the functional consequences of *BIN1* deletion at the single neuronal level (Fig. 6A). We identified a similar number of single units per recording electrode between genotypes (WT: 4.92±2.34; KO: 5.27±2.45), indicating that *BIN1* deletion does not affect the density of active neurons within culture. However, we observed reduced single-unit activity (SUA) frequency (Fig. 6B) and higher SUA amplitude (Fig. 6C) in *BIN1* KO compared to WT ASCL1-hiNs. Interestingly, we could not detect significant changes in the number of bursts per neuron (WT: 11.01±6.71; KO: 10.36±8.59), although both the burst duration and the number of spikes within a burst were significantly lower in *BIN1* KO compared to WT ASCL1-hiNs (Fig. 6D-E). We also observed a prominent temporal disorganization of *BIN1* KO hiNs activity by computing the array-wide spike detection rate (ASDR, Fig. 6G), which reveals the strength of the synchronized population activity, and the autocorrelograms of SUAs (Fig. 6H-I), which allows the apprehension of synchronized periodicity. These analyses revealed that most spikes of *BIN1* WT neurons were organized in bursts occurring at periodic intervals of about 8-10 s, whereas the spikes of *BIN1* KO neurons were randomly distributed, leading to a higher percentage of spikes occurring outside of bursts compared to WT neurons (Fig. 6J).

**Figure 5:**
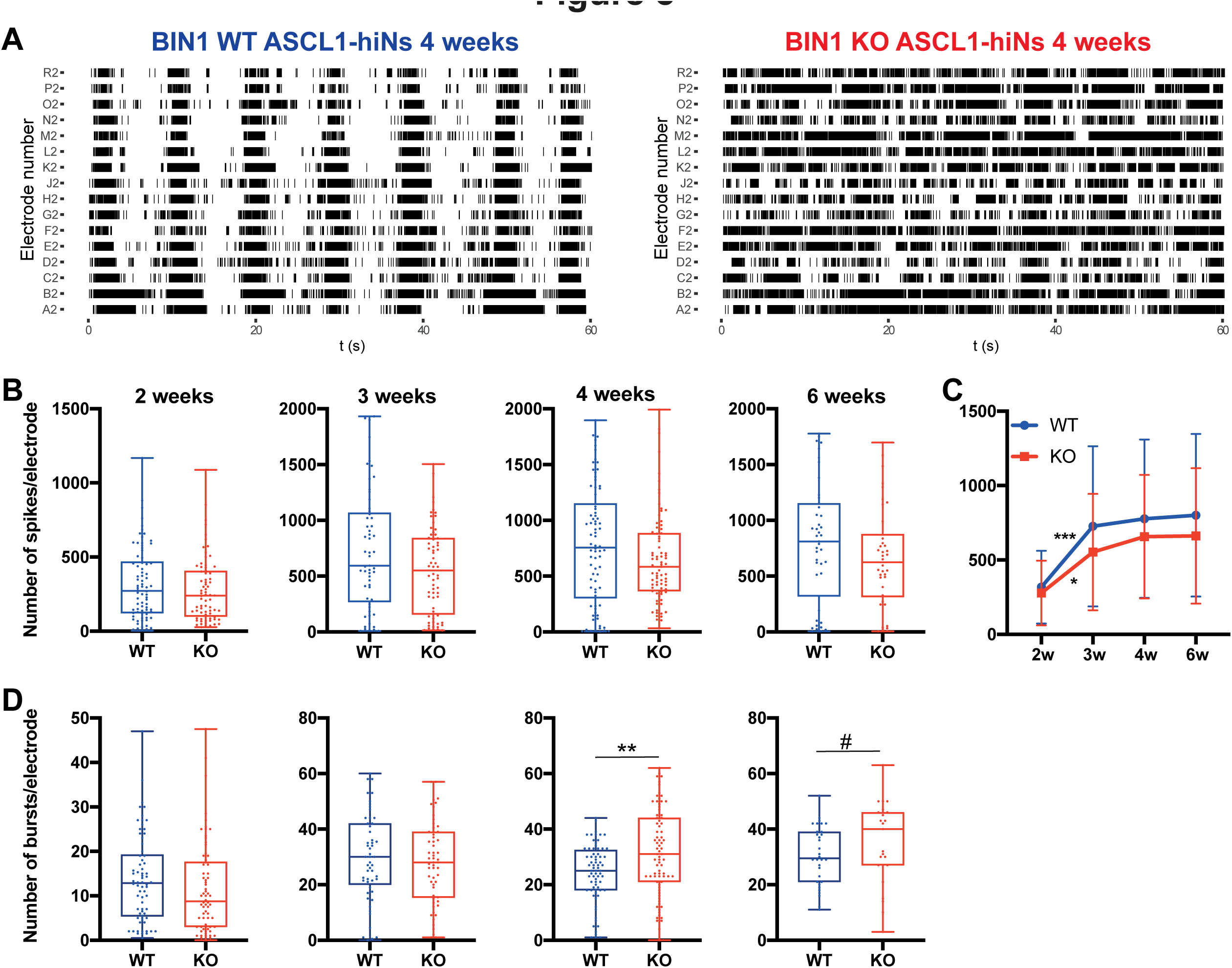
BIN1 KO ASCL-hiNs have a higher spike burst frequency compared to WT. (A) Raster plots showing MUA recorded for 1 minute in *BIN1* WT and KO ASCL1-hiNs after 4 weeks of differentiation. Each line represents one electrode localized side-by-side in our microfluidic/MEA array (as in panel A). (B-C) Quantification of the number of detected spikes at different time points (*p=0.0141; ***p=0.0006; Two-way ANOVA followed by Tukey’s multiple-comparison test; n= 5 for each genotype). (D) Quantification of the number of spike bursts at different time points (**p=0.004; ^#^p=0.0888; Mann-Whitney test).

**Figure 6:**
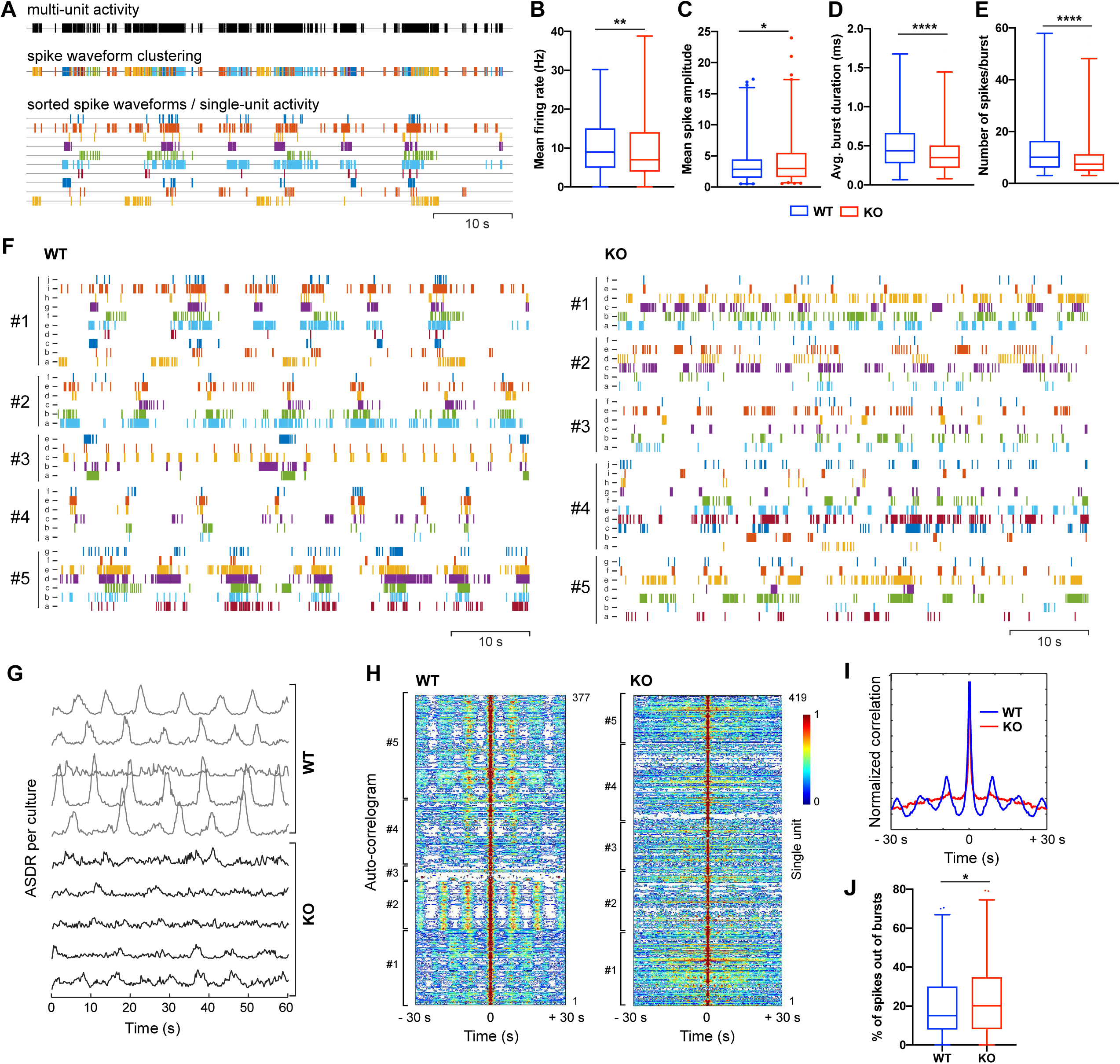
Disorganization of neuronal activity in BIN1 KO ASCL1-hiNs. (A) Raster plots showing the decomposition of multi-unity activity (MUA, black lines) into single-unit activity (SUA, colored lines) using spike waveform clustering. (B-E) Quantification of single-neuron firing rate (B; **p=0.0034), spike amplitude (C; *p=0.0106), burst duration (D; ****p<0.0001) and number of spikes per burst (E; ****p<0.0001) at 4 weeks. Mann-Whitney test; n= 5 independent experiments; WT: 376 neurons; KO: 416 neurons). (F) Raster plots showing SUA recorded from 5 different electrodes of *BIN1* WT (left) or KO (right) ASCL-hiNs cultures after 4 weeks of differentiation. (G) Array-wide spike detection rate (ASDR) plots based on SUA recorded in *BIN1* WT and KO ASCL1-hiNs cultures. Each line represents one independent culture batch. (H-I) Normalized autocorrelogram heatmap (H, each line refers to one SUA) and averaged correlation (I) for all SUAs recorded in 5 independent *BIN1* WT and KO ASCL1-hiNs cultures. (J) Percentage of spikes outside of bursts (*p=0.0417, Mann-Whitney test).

### Altered electrical activity of BIN 1KO ASCL1-hiNs is associated with normal synapse numbers and altered TAU phosphorylation

These changes in neural network activity observed in *BIN1* KO hiNs could be explained, among other things, by a reduced synaptic connectivity, as observed in our long-term COs cultures (Fig. 3). To test this possibility, we first quantified the number of synaptic contacts in *BIN1* WT and KO ASCL1-hiNs cultures. In contrast with COs, we did not detect any significant differences in the number of putative synaptic contacts (% SYP assigned) in *BIN1* KO compared to WT ASCL1-hiNs, neither after 4 nor 6 weeks of differentiation (Fig. 7A-D). Next, we quantified the number and activity of glutamatergic synapses by using real-time imaging of ASCL1-hiNs expressing the glutamate sensor iGLUSnFr [30]. In accordance with our observations based on immunocytochemistry, we did not detect differences neither in the number of glutamatergic synapses (active spots) nor in the frequency of events (change in fluorescence levels in active spots) in *BIN1* KO compared to WT ASCL1-hiNs (Sup. Fig. 3; Sup. Movies 1 and 2), indicating that changes in neuronal activity observed in our cultures are not related to changes in synaptic transmission.

**Figure 7:**
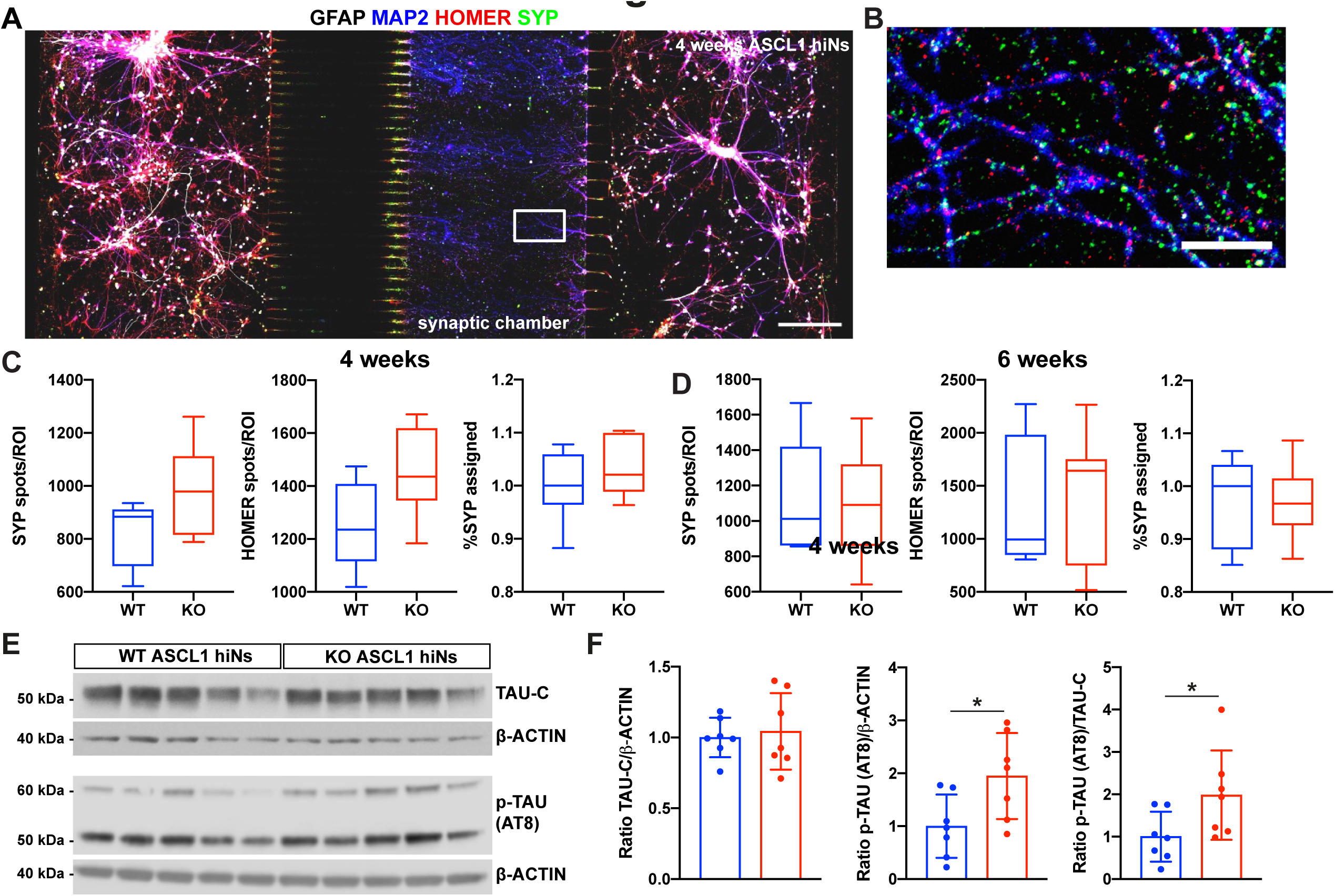
Similar synaptic density and higher TAU phosphorylation in BIN1 K O compared to WT ASCL1-hiNs. (A-B) Immunocytochemistry using the astrocyte marker GFAP, neuronal marker MAP2, pre-synaptic marker SYP and post-synaptic marker HOMER1 in *BIN1* WT ASCL1-hiNs after 4 weeks of differentiation in a three-chamber microfluidic device. Scale bar = 200 μm. Rectangular box in A is magnified in B, allowing the identification of putative synaptic contacts. (C-D) Fraction of SYP spots assigned by HOMER1 spots in MAP2 processes at 4 (C) and 6 weeks (D) ASCL1-hiNs cultures (n= 8 independent devices for each genotype). (E) Western blot for total TAU protein C-terminal (TAU-C), phosphorylated (p)-TAU at Ser202, Thr205 (AT8) and β-ACTIN in 4-week-old ASCL1-hiNs cultures. (F) Quantification of TAU-C/β-ACTIN and p-TAU/TAU-C levels in BIN1 KO ASCL1-hiNs normalized to WT (*p=0.0262; Mann-Whitney test).

Taking advantage of this culture system comprising enriched neuronal populations, we also sought to confirm whether BIN1 deletion could be associated with changes in APP processing or Tau phosphorylation. To this end, we measured the extracellular levels of amyloid-beta (Aβ) peptides, as well as the intracellular levels of full-length APP and APP β-CTF, total and phosphorylated TAU proteins in ASCL1-hiNs cultures. Like COs, we did not detect any significant differences neither in the extracellular levels of Aβ1-x or Aβ1-42, nor in the intracellular levels of APP or APP β-CTF in *BIN1* KO compared to WT ASCL1-hiNs (Sup. Fig. 4). However, in contrast with our observations in COs, we observed significantly higher levels of phospho-TAU (Ser202, Thr205) relative to β-ACTIN and total TAU in in *BIN1* KO compared to WT ASCL1-hiNs (Fig. 7E-F). Together, these observations may suggest that *BIN1* deletion primarily impairs neuronal intrinsic properties regulating electrical activity and Tau phosphorylation prior to detectable changes in synaptic communication (observed only in long-term cultures) and independently of alterations in APP processing.

### BIN1 regulates neuronal Ca**^2+^** dynamics through LVGCCs

In neurons, electrical activity is always accompanied by an influx of Ca2+ ions, which play a fundamental role in the regulation of neuronal firing and activity-dependent gene transcription [31]. We therefore postulated that reduced BIN1 expression in human glutamatergic neurons could affect Ca2+ dynamics, as previously suggested for cardiomyocytes [32]. To directly test this possibility, we first studied Ca^2+^ dynamics in *BIN1* WT and KO ASCL1-hiNs using real-time calcium imaging experiments. We observed spontaneous synchronous calcium transients among adjacent cells both in *BIN1* WT and KO ASCL1-hiNs cultures (Sup. Movies 3 and 4). By quantifying calcium spike transients (> 2 standard deviations above the noise level) we showed a significantly higher frequency of Ca^2+^ transients in *BIN1* KO compared to WT ASCL1-hiNs (Fig. 8A,B and D). Moreover, the dynamics of individual Ca^2+^ transients in *BIN1* KO were qualitatively different from WT ASCL1-hiNs (Fig. 8C). These differences could be quantitatively measured by a longer time to reach the maximum intracellular Ca^2+^ levels and to recover baseline levels (Fig. 8E-F).

**Figure 8:**
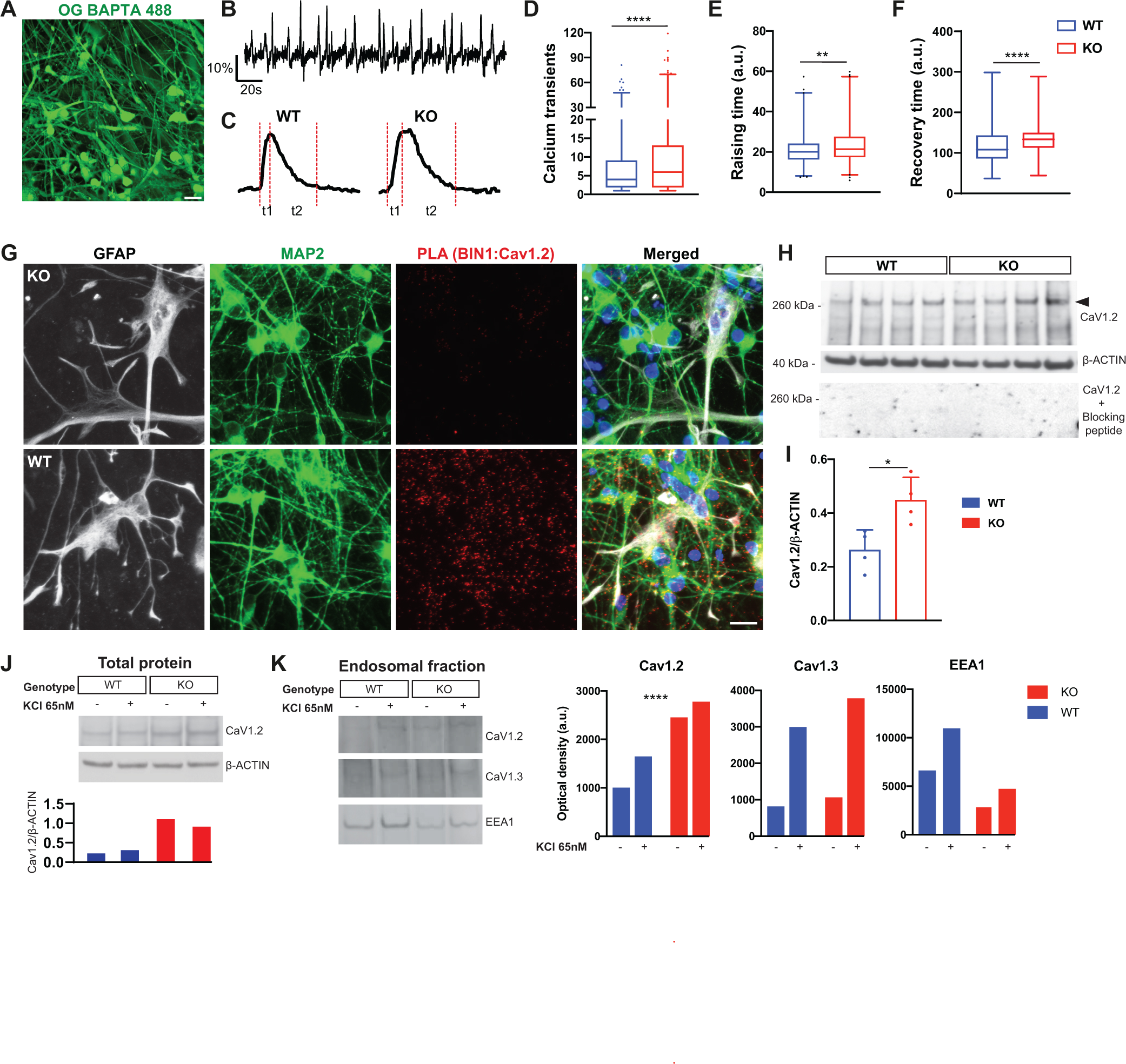
Altered frequency of calcium transients in BIN1 KO ASCL1-hiNs. (A) Snapshot of a 4-week-old ASCL1-hiNs culture labeled with Oregon green BAPTA. (B) Representative plot of fluorescence change over time in 1000 frames. (C) Representative traces showing the fluorescence changes in *BIN1* WT and KO ASCL1-hiNs. Red dashed lines indicate the time to reach the fluorescence maximal intensity (raising time - t1) and to return to baseline (recovery time - t2). (D) Quantification of calcium transients in *BIN1* WT and KO ASCL1-hiNs (****p<0.0001; Mann-Whitney test; n= 3 independent cultures for each genotype; number of active cells per condition: 754 (WT), 1006 (KO)). (E-F) Quantification of rising time (t1) and recovery time (t2) for calcium transients (**p=0.0022; ****p<0.0001; Mann-Whitney test). (G) Images showing PLA spots using anti-BIN1 and anti-Cav_1.2_ antibodies in 4-week-old *BIN1* WT and KO hiNs. Cells were also immunolabeled for the neuronal marker MAP2 (green), the astrocyte marker GFAP (white), and stained with DAPI (blue). (H) Western blot for Cav1.2 (without and with blocking peptide) and β-ACTIN in 4-week-old ASCL1-hiNs cultures. (I) Quantification of Cav1.2/β-ACTIN levels in *BIN1* WT and KO ASCL1-hiNs cultures (*p=0.0286; Unpaired t-test). (J) Western blot for Cav_1.2_ and β-ACTIN in the total protein extracts from 4-week-old ASCL1-hiNs treated with KCl (+) or vehicle (-). Plot shows the quantification of Cav_1.2_ normalized by β-ACTIN. (K) Western blot for Cav_1.2_, Cav_1.3_ and EEA1 in the endosomal protein extracts from 4-week-old ASCL1-hiNs treated with KCl (+) or vehicle (-). Plot shows the optical density of these proteins (****p<0.0001; Chi-square test).

In human heart failure, BIN1 expression is reduced, leading to an impairment in Cav1.2 trafficking, calcium transients, and contractility (Hong et al., 2012). Thus, we sought to determine if BIN1 could also interact and regulate LVGCC expression in human neurons. To that, we performed proximity ligation assay (PLA) to probe a possible interaction between BIN1 and Cav1_.2_ or Cav_1.3_, the two LVGCCs expressed in ASCL1-hiNs (Sup. Fig. 5). We observed a widespread BIN1-Cav_1.2_ PLA signal (Fig. 8G) and, to a lesser extent, a BIN1-Cav_1.3_ PLA signals in neurons (Sup. Fig. 5). Next, we quantified neuronal LVGCC protein by western blotting and observed higher total Cav_1.2_ expression in *BIN1* KO compared to WT ASCL1-hiNs (Fig. 8H-I). Protein expression of neither Cav_1.3_, nor the members of the Cav_2_ family (Cav_2.1_, Cav_2.2_ and Cav_2.3_) were altered in the same cultures (Sup. Fig. 5), suggesting a specific regulation of Cav_1.2_ expression by BIN1.

Notably, LVGCCs are key regulators of neuronal firing [29] and activity-dependent internalization of these channels is a key mechanism in firing homeostasis [33]. We thus set out to investigate whether *BIN1* deletion could impair this mechanism in human neurons. We stimulated ASCL1-hiNs with KCl 65nM for 30 min to induce neuronal depolarization and collected total and endosomal proteins for analysis. We confirmed a higher global level of Cav_1.2_ in *BIN1* KO compared to WT ASCL1-hiNs that was independent of KCl treatment (Fig. 8J). However, Cav_1.2_ expression in the endosomal fraction was 50% higher after KCl treatment in *BIN1* WT, whereas this rise was only of 10% in *BIN1* KO ASCL1-hiNs (Fig. 8K). This effect was specific for Cav_1.2_ since both early endosome antigen 1 (EEA1) and Cav_1.3_ expression rose in both *BIN1* WT and KO ASCL1-hiNs at similar levels after KCl treatment (Fig. 8K). Altogether, these results indicate that BIN1 regulates activity-dependent internalization and expression of Cav_1.2_ in human neurons.

### Treatment with the calcium channel blocker nifedipine partly rescues electrical and gene expression alterations in BIN1 KO ASCL1-hiNs

To investigate whether the network dysfunctions observed in *BIN1* KO ASCL1-hiNs may be related to the higher expression of Cav_1.2_, we treated these cells with a physiologically relevant concentration (50nM) of the Cav_1.2_ blocker nifedipine [34]) for 2 weeks and recorded neuronal activity using MEA electrophysiology. We observed a partial recovery of the oscillatory pattern of neuronal electrical activity observed in WT cells (Fig. 9A). Interestingly, the percentage of spikes outside bursts was not affected by nifedipine treatment in *BIN1* WT but was significantly lower in *BIN1* KO ASCL1-hiNs (Fig. 9B), indicating a partial recovery of burst organization. To note, no difference in firing rates was observed whatever the models and conditions (Fig. 9C). After 2 weeks of nifedipine treatment (4 weeks of differentiation), we also performed snRNA-seq experiments and recovered a total of 1537 cells (n= 2 independent culture batches), which were mapped into the 7 clusters described before (Fig. 4; Sup. Figure 6). Using Wilcoxon test, we found that nifedipine treatment down-regulated several genes in *BIN1* KO ASCL1-hiNs, especially in the GluNeu-II population (Sup. Table 6). Comparison of GO term enrichments between nifedipine-treated and untreated *BIN1* KO vs WT GluNeu-II population revealed a consistent reduction of the enrichment for several terms associated with ion channel activity and synapse transmission in nifedipine-treated BIN1 KO cells (Figure 9D; Sup. Table 7). Altogether, these data support the view that BIN1 contributes to the regulation of electrical activity and gene expression through the regulation of Cav_1.2_ expression/localization in human neurons.

**Figure 9:**
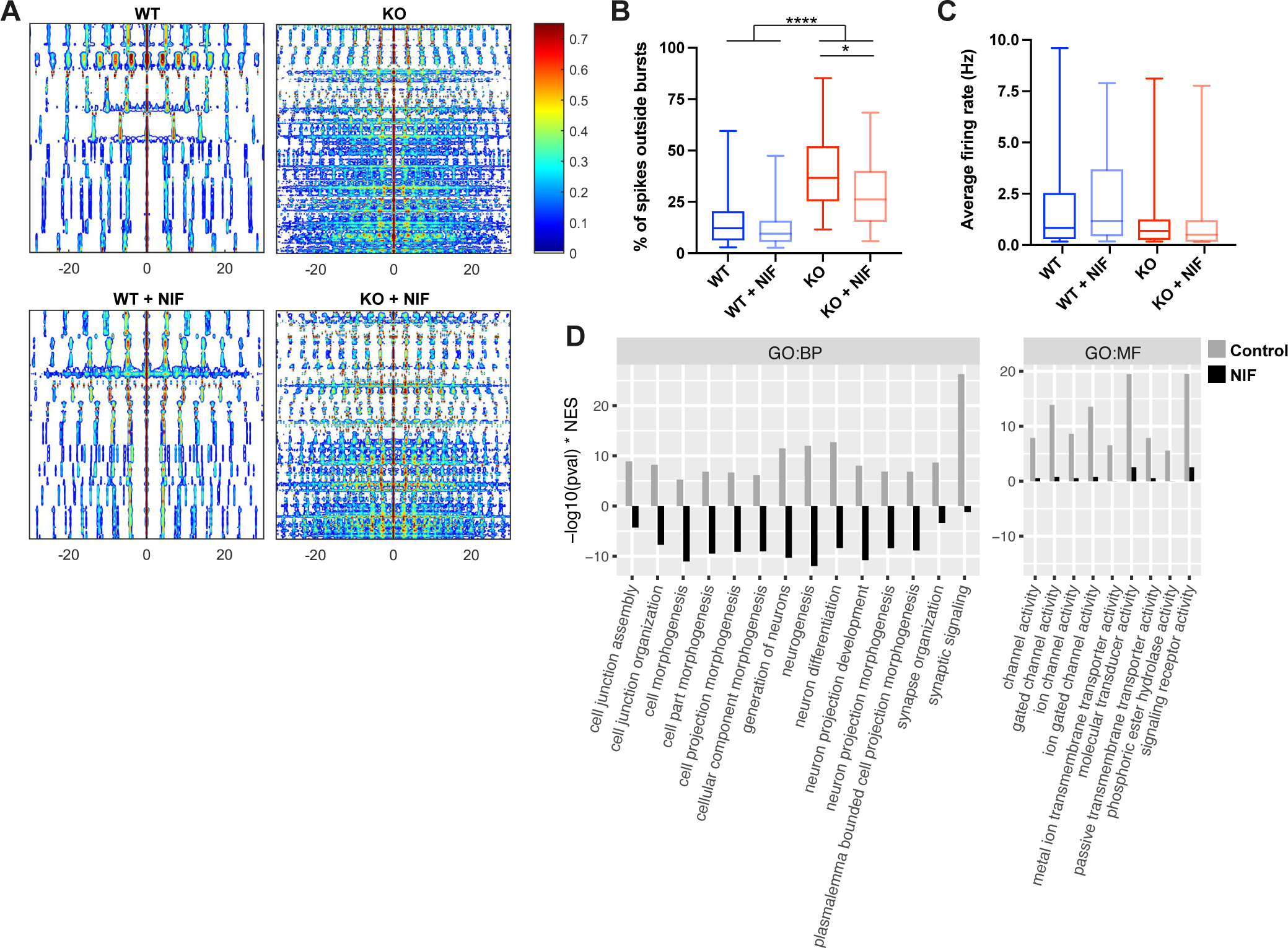
Nifedipine treatment partly rescues electrical and gene expression alteration in BIN1 KO hiNs. (A) Auto-correlograms of 4-week-old *BIN1* WT and KO hiNs treated or not with 50 nM Nifedipine for 2 weeks. (B) Percentage of spikes outside of bursts (WT vs WT+NIF: **p_adj_=0.0034; WT vs KO: *p_adj_=0.0124; Dunn’s multiple comparison test). (C) Average firing rates. (D) Comparison of the enrichment for GO terms in genes upregulated in KO vs WT and KO + nifedipine (NIF) vs WT GluNeu-II using fast gene set enrichment analysis (FGSEA). The FGSEA results are shown for the top15 GO terms with reduce enrichment in KO NIF.

### Molecular alterations in BIN1 KO organoids and ASCL-hiNshiNs are also present in glutamatergic neurons of the human brain at late stages of AD pathology

We finally sought to evaluate whether molecular alterations in our neural models may recapitulate some of those observed in the brain of AD cases. For this purpose, we used a publicly available snRNA-seq dataset generated from the entorhinal cortex (EC) and superior frontal gyrus (SFG) of AD patients at different Braak stages [35]. We first observed a progressive and significant decrease in BIN1 mRNA levels in glutamatergic neurons (Fig. 10A), suggesting that reduced BIN1 expression in this cell type may be a common feature occurring in the AD pathology progression. We then compared DEGs identified in BIN1 KO glutamatergic neurons (either from COs or ASCL1-hiNs) with those identified in the same cell subtype of AD brains (Sup. Table 8). Remarkably, DEGs identified in *BIN1* KO glutamatergic neurons (either from COs or ASCL1-hiNs) showed a statistically significant overlap with DEGs detected in this cell population in AD brains at different Braak stages (Fig. 10B). In astrocytes, however, a similar significant overlap could only be observed between COs and AD brains (Fig. 10B). GO analysis based on DEG overlap between *BIN1* KO ASCL1-hiNs and AD brain glutamatergic neurons indicated significant enrichment for pathways associated with glutamate receptor activity and gated channel activity (Fig. 10C; Sup. Table 7). Similarly, DEG overlap between *BIN1* KO COs and AD brain glutamatergic neurons was significantly enriched for genes associated with glutamate receptor activity, gated channel activity and calcium ion binding (Fig. 10D; Sup. Table 7). No significant enrichment was observed for DEG overlap between *BIN1* KO COs and AD brain astrocytes (data not shown).

**Figure 10:**
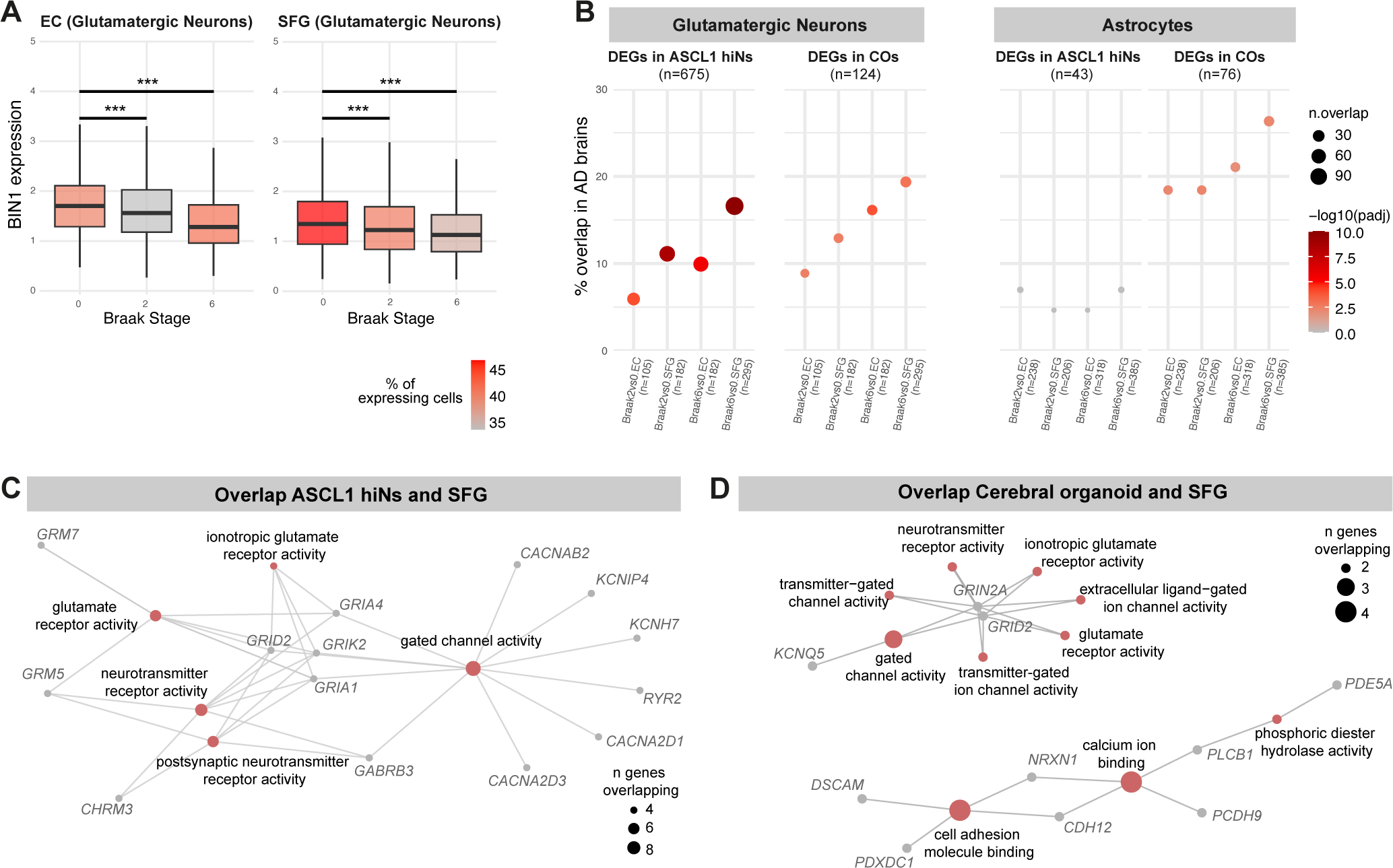
Similar molecular alterations in BIN1 KO hiNs and glutamatergic neurons of the AD brain. (A) Box plot representing *BIN1* mRNA in expression through different Braak stages in entorhinal cortex (EC) and superior frontal gyrus (SFG) (***p_adj_<0.001; Wilcoxon test). (B) Dot plot representing the overlap between DEGs identified in glutamatergic neurons of the AD brain and *BIN1* KO ASCL1-hiN cultures (left) or *BIN1* KO COs (right). (C-D) Network representation of enriched GO terms in overlapping DEGs between AD brains and glutamatergic neurons in culture. Enriched GO terms were identified using over-representation test.

## Discussion

In this work, we show that the AD genetic risk factor *BIN1*, plays a critical role in the regulation of neuronal firing homeostasis and gene expression in glutamatergic neurons. Complete deletion of *BIN1* gene in these neurons is sufficient to alter the expression of the LVGCC Cav_1.2_, leading to altered calcium homeostasis and neural network dysfunctions in human neurons *in vitro* . These functional changes are correlated with changes in the expression of genes involved in synaptic transmission and ion transport across the membrane, as well as elevated Tau phosphorylation. In long-term neuronal cultures using COs, we show that reduced BIN1 expression is associated with fewer synapses and specific gene expression alterations in glutamatergic neurons associated with activity-dependent transcription. Notably, reduced BIN1 expression in human-induced glutamatergic neurons is sufficient to elicit gene expression alterations that are also present in AD and converge to biological processes related with calcium homeostasis. Together, our findings support the view that altered BIN1 expression in glutamatergic neurons may contribute to AD pathophysiology by dysregulating neuronal firing homeostasis via LVGCCs.

Neuronal network dysfunctions are observed in AD patients at early stages of the disease and precede or coincide with cognitive decline [36–38]. Under physiological conditions, neuronal networks can maintain optimal output through regulation of synaptic plasticity and firing rate [39]. Our results suggest that normal levels of BIN1 expression in glutamatergic neurons are fundamental to regulating neuronal firing rate homeostasis. Accordingly, *BIN1* deletion in hiNs is sufficient to dysregulate network oscillations even without impacting the number of functional synaptic contacts, suggesting that the desynchronization observed in *BIN1* KO hiNs circuits are a consequence of disordered homeostatic controls of neuronal activity. At later time-points (COs), glutamatergic neurons show both gene expression alterations indicative of altered electrical activity and reduced synaptic densities, which could indicate a synaptic down-scaling in response to earlier augmented electrical activity (Dörrbaum et al., 2020).

One key mechanism controlling neuronal spiking activity is the regulation of Ca^2+^ homeostasis [29,31,40]. Boosted neuronal electrical activity induces the turnover of LVGCCs from the plasma membrane through endocytosis [33] and regulates the transcription of genes encoding for calcium-binding proteins and calcium-mediated signaling [41], mechanisms aiming to restore local Ca^2+^ signaling cascades and protect cells against aberrant Ca^2+^ influx. We show that BIN1 interacts with Cav_1.2_ in hiNs, similar to previous findings in cardiac T tubules [32] and provide evidence supporting a novel role for BIN1 in the regulation of activity-dependent internalization of Cav_1.2_ in human neurons, thus linking the known role of BIN1 in endocytosis [12] to firing homeostasis in human neurons via the LVGCC. These results confirm in human neurons the interaction between endogenous neuronal BIN1 and Cav_1.2_ as previously suggested in mouse hippocampal neurons, overexpressing a Bin1-mKate fused protein [18].

Loss of Ca^2+^ homeostasis is an important feature of many neurological diseases and has been extensively described in AD [42,43]. Interestingly, DEGs identified in *BIN1* KO glutamatergic neurons in our different cell culture models are enriched for calcium-related biological processes. This is also observed for DEGs detected in glutamatergic neurons of the AD brain at late stages of pathology when the expression levels of BIN1 in those cells is also decreased. Thus, it is plausible to speculate that reduced expression of *BIN1* in glutamatergic neurons may contribute to the breakdown of Ca^2+^ homeostasis in the AD brain, potentially contributing to neuronal circuit dysfunctions. Consistent with this hypothesis, we have previously shown a significant reduction in the expression of the transcript encoding for the neuron-specific BIN1 isoform 1 in bulk RNA-sequencing data from a large number of AD patients [7] and we show in this work that *BIN1* expression is reduced in glutamatergic neurons of AD brains at late Braak stages. Moreover, it has been recently shown that Bin1 conditional knockout in neurons and glial cells of the mouse forebrain is sufficient to elicit gene expression changes associated with calcium-dependent mechanisms (Ponnusamy et al., 2022), further supporting the interpretation that BIN1 plays an important role in cellular processes involved in Ca^2+^ homeostasis in the brain.

Lastly, we show that treatment with the clinically approved calcium channel blocker nifedipine for only 2 weeks is sufficient to partly recover electrical and gene expression alterations in *BIN1* KO hiNs. These findings further support our interpretation that changes in gene expression and electrical activity observed in *BIN1* HET and KO hiNs are a direct consequence of reduced BIN1 expression in glutamatergic neurons and not a possible artifact of CRISPR/Cas9 gene-editing in our hiPSC lines. Moreover, together with our observations of reduced BIN1 expression and transcriptional alterations affecting biological processes related to calcium homeostasis in human-induced glutamatergic neurons of the human brain at late stages of AD pathology, our data strongly support a link between BIN1 and calcium homeostasis.

Thus, it is plausible to speculate that *BIN1* depletion in glutamatergic neurons primarily undermines Ca^2+^ homeostasis, leading to changes in neuronal electrical activity. In a later stage, gene expression and circuit-level alterations such as synapse loss would occur, likely because of altered neuronal electrical activity. A corollary to this model would be that early treatments aiming to restore Ca^2+^ homeostasis and neuronal electrical activity may have a beneficial impact in AD onset and progression. Supporting this notion, both a Mendelian randomization and a retrospective population-based cohort study found evidence suggesting that treatment with Ca^2+^ channel blockers in human patients are associated with a reduced risk of AD [44,45]. In the future, it would be interesting to study the impact of these drugs for AD onset/progress as a function of genetic variants in the *BIN1* locus.

An important limitation of our work is the absence of microglial cells in our models, hampering the study of BIN1 roles in this cell type of high relevance to AD pathology [46]. However, our findings provide fundamental information about the molecular and cellular processes impacted by reduced BIN1 expression in human neurons, which will require further confirmation in the brain of patients carrying AD-related *BIN1* genetic variants. Moreover, we studied the impact of heterozygous and knockout mutations, which do not necessarily represent the consequences of AD-related *BIN1* genetic variants in gene expression. Nevertheless, it is tempting to speculate that slight changes in *BIN1* expression provoked by those variants could progressively deteriorate neuronal functions in the human brain, contributing to AD pathogenesis in the elderly brain.

## Declarations

### Ethics approval and consent to participate

The datasets used and/or analysed during the current study are available from the corresponding author on reasonable request.

**Consent for publication** Not applicable.

### Availability of data and materials

Not applicable.

## Funding

This work was co-funded by the European Union under the European Regional Development Fund (ERDF) and by the Hauts de France Regional Council (contract no.18006176), the MEL (contract_18006176), and the French State (contract no. 2018-3-CTRL_IPL_Phase2). This work was partly supported by the French RENATECH network (P-18-02737), Fondation pour la recherche médicale (ALZ201912009628, ALZ201906008477), PTR-MIAD, Fondation Recherche Alzheimer and by the Sanofi i-Awards Europe 2019. This work was also funded by the Lille Métropole Communauté Urbaine and the French government’s LABEX DISTALZ program (Development of innovative strategies for a transdisciplinary approach to Alzheimer’s disease). The UMR 8199 LIGAN-PM Genomics platform (Lille, France) belongs to the ’Federation de Recherche’ 3508 Labex EGID (European Genomics Institute for Diabetes; ANR-10-LABX-46) and was supported by the ANR Equipex 2010 session (ANR-10-EQPX-07-01; ’LIGAN-PM’). The LIGAN-PM Genomics platform (Lille, France) is also supported by the FEDER and the Region Nord-Pas-de-Calais-Picardie and is a member of the “France Génomique” consortium (ANR-10-INBS-009).

### Authors’ contributions

Conceptualization, M.R.C.; Methodology, M.R.C., F.D., D.K., C.M.Q.; Investigation, M.R.C., O.S., B.S.L., A.R.M.F., J.G., A.C., A.P., D.S.W., F.D., K.G., D.K., A.C.V., C.M.Q.; Writing -Original Draft, M.R.C.; Writing - Reviews & Editing, M.R.C., F.D., J.C.L., C.M.Q., D.K.; Figures preparation: M.R.C., F.D., A.P., B.S.L., A.R.M.F., D.K. Supervision, M.R.C., F.D., J.C.L., D.K.; Funding Acquisition, M.R.C., J.C.L., F.D, D.K., P.A., A.B. All authors have read and approved the manuscript.

## Acknowledgements

The authors thank the BICeL platform of the Institut Biologie de Lille and the Vect’UB viral platform (INSERM US 005 – CNRS 3427 – TBMCore, Université de Bordeaux, France). The authors thank Karine Blary at the IEMN Lille for the microfabrication work. The Maestro Pro multiwell microelectrode array was acquired with the “Prix Claude Pompidou pour la Recherche sur l’Alzheimer (2021)” to MRC.

## Figure legends

### Supplementary data

**Sup. Movies 1 and 2:** Time-series of 1000 frames taken from *BIN1* WT and KO ASCL1-hiNs transduced with iGLUSnFr after 2 weeks of differentiation and imaged 2 weeks later. Videos are played at 100 fps.

**Sup. Movies 3 and 4:** Time-series of 1000 frames taken from *BIN1* WT and KO ASCL1-hiNs after 4 weeks of differentiation and labeled with Oregon Green BAPTA and imaged. Videos are played at 100 frames per second (fps).

**Sup. Table 1**: DEGs identified in different cell types/subtypes of COs.

**Sup. Table 2**: GO terms enriched for DEGs identified in different cell types/subtypes of COs.

**Sup. Table 3**: List of ARGs used for hypergeometric test in Cell-ID.

**Sup. Table 4**: DEGs identified in different cell types/subtypes of ASCL1-hiNs cultures.

**Sup. Table 5**: GO terms enriched for DEGs identified in different cell types/subtypes of ASCL1-hiNs cultures.

**Sup. Table 6**: DEGs identified in different cell types/subtypes of ASCL1-hiNs cultures treated with nifedipine.

**Sup. Table 7**: GO terms enriched for DEGs identified in different cell types/subtypes of ASCL1-hiNs cultures treated with nifedipine.

**Sup. Table 8**: DEGs identified in different cell types/subtypes of the AD brain.

**Sup. Table 9**: GO terms enriched for DEGs commonly identified in *BIN1* HET or KO cells and the AD brain.

## Materials and methods

### Maintenance of isogenic hiPSC lines and neural induction

Isogenic hiPSCs (ASE 9109, Applied StemCell Inc. CA, USA) modified for BIN1 in exon 3 were generated by CRISPR/Cas9. Homozygous null mutants for BIN1 had a 5 bp deletion on one allele and an 8 bp deletion on the other allele. Heterozygous for BIN1 had a 1bp insertion on one allele. All hiPSCs, and all subsequent human induced neural progenitor cells (hiNPCs), hiNs, human induced astrocytes (hiAs), and cerebral organoids derived thereof, were maintained in media from Stemcell Technologies, Vancouver, Canada. Maintenance of cell cultures and organoids were done in adherence with manufacturer’s protocols which can be found on the webpage of Stemcell Technologies. hiPSCs were maintained in mTeSR1 medium in non-treated cell culture dishes/plates pre-coated with vitronectin. Cell numbers and viability were recorded using a LUNA™ Automated Cell Counter.

To generate BIN1 WT and KO hiNPCs, we used the dual SMAD-inhibition (SMADi) method (Chambers et al., 2009). Briefly, hiPSCs are cultured and passaged for 3 times in STEMdiff™ Neural Induction Medium + SMADi (Stemcell Technologies) for up to 3 weeks. After this period, hiNPCS are maintained for up to 10 passages in treated cell culture dishes pre-coated with poly-L-ornithine (PLO) and laminin (5 µg/mL), using STEMdiff™ Neural Progenitor Medium (NPM; Stemcell Technologies). PLO solution was made in water (0.001%) while laminin was diluted in PBS with Ca2+ and Mg2+.

### Generation of Cerebral Organoids

Cerebral organoids (3D Cultures) were generated from BIN1 WT, HET and KO hiPSCs using a 4-stage protocol (Lancaster et al., 2013). The first step was the Embryoid Body (EB) Formation Stage, where hiPCSc at 80%-90% confluency were detached from the Vitronectin XF substrate using Accutase (#AT-104, Innovative Cell Technologies). To form the EB, 9000 cells were plated per well in a 96-well round-bottom ultra-low attachment plate containing EB seeding medium (Stem Cell Technologies). At day 2 and day 4, EB formation media is added in the wells. After five days, the EBs were transferred to a 24-well ultra-low attachment plate containing Induction Medium (Stem Cell Technologies), where each well receives 1-2 EBs. Two days later, the EBs were ready for the Expansion Stage. The EBs were embedded in Matrigel (Corning) and transferred to a 24-well ultra-low adherent plate with Expansion Medium (Stem Cell Technologies). After three days, the medium was replaced by Maturation Medium (Stem Cell Technologies) and the plate was placed in an orbital shaker (100 rpm speed). During this final Maturation Phase, 75% medium change was done on a biweekly basis. Organoids were allowed to mature for a period of 6.5 months.

### Generation of pure human neuronal cultures from hiNPCs

We differentiated neurons from virus-transduced hiNPCs according to an adapted protocol (Zhang et al., 2013; Yang et al., 2017). Briefly, hiNPCs are first virally-transduced with the TTA lentiviral construct and a passage later, the TetO-Ascl1-Puro lentiviral construct was transduced. These cells are maintained in NPM medium and expanded prior to differentiation. For differentiation of human induced neurons (hiNs), hiNPCs are plated onto PLO/laminin-coated imaging plates at density 100,000 cells/well in NPM. After 24 h, complete BrainPhys medium (StemCell Technologies) is added 1:1 together with 2 µg/mL doxycycline (Sigma-Aldrich) to induce TetO gene expression. The following day, 1 µg/mL puromycin (Sigma-Aldrich) was added to start cell selection. After 2-3 days (depending on the efficiency of antibiotic selection), 50,000 human cortical astrocytes were added in each well with BrainPhys containing doxycycline. After 24 h, 2 µM of Ara-C (Cytosine β-D-arabinofuranoside; Sigma-Aldrich) was added to arrest the proliferation of astrocytes. Half of the medium in each well was changed biweekly with fresh BrainPhys medium containing doxycycline until the 14^th^ day. After that, the biweekly medium change was performed only with BrainPhys. Differentiation was allowed to continue for another 2-4 weeks prior to subjecting the cells to various experimental manipulations. Human cortical astrocytes (HA; ScienCell # 1800) were sourced from ScienCell Research Laboratories, CA, USA and grown up ^2^to 10 passages on poly-l-lysine coated 75 cm flasks using astrocyte medium (ScienCell # 1801) supplemented with astrocyte growth supplement (AGS, ScienCell #1852) and 10 ml of fetal bovine serum (ScienCell # 1800).

This co-culture system was characterized using snRNA-seq after 4 weeks of differentiation showing that 70% of cells (n=7120 from 5 independent culture batches) expressed the pan-neuronal markers SOX11, SNAP25, DCX and RBFOX3, with 66% of cells co-expressing the glutamatergic neuron marker SLC17A6, less than 1.5% of cells co-expressing the GABAergic neuron markers DLX1, GAD1 and GAD2, and 5% of cells co-expressing low levels of markers of both neuronal subtypes. The remaining cells, immature astrocytes (Astro-I), mature astrocytes (Astro-II) and undifferentiated NPCs, represented about 15%, 8%, 4% of the cells, respectively. The first two cell populations likely represent two different states of astrocytes added to the cultures, whereas NPCs are likely cells that failed to reprogram into hiNs despite ASCL1 transduction. Thus, neuronal classes generated in this system comprise a large fraction of glutamatergic neurons (84%), a small proportion of GABAergic neurons (2.5%) and unspecified neurons co-expressing low levels of markers of both neuronal subtypes (13.5%).

### Culture of hiNs in Microfluidic Devices

Three-compartment microfluidic neuron culture devices were used in which the presynaptic and postsynaptic chambers are connected to the synaptic chamber by respectively long and short micro-channels. Details of the microfluidic device design and fabrication have been previously described (Kilinc et al, 2020).

The homemade devices were placed individually in Petri dishes for easy handling and UV sterilized for 30 min before coating for cell adhesion. The primary surface coating consisted of poly-L-lysine (Sigma-Aldrich) at 20 µg/mL in borate buffer (0.31% boric acid, 0.475% sodium tetraborate, pH 8.5). All coated devices were incubated overnight at 37°C, 5% CO2. After a wash with DPBS, devices were then coated with 20 μg/mL laminin in DPBS and incubated overnight at 37°C in 5% CO2. The following day, devices were carefully washed once with DPBS before cell plating.

In total, 30,000 hiNPCs resuspended in NPM containing 10 µM of Y-27632 ROCK inhibitor were seeded per device, half at the entrance of the presynaptic somatic chamber and half at the entrance of the postsynaptic somatic chamber. Microfluidic devices were microscopically checked at the phase contrast to ensure the cells were correctly flowing into chambers. After a minimum of 5 minutes to allow the cells to attach, devices were filled with NPM (containing 10 µM of Y-27632 ROCK Inhibitor). Water was added to the Petri dishes to prevent media evaporation, and these were then incubated at 37°C in a humidified 5% CO2 incubator. For induced neuron culture from hiNPCs transduced for Ascl1, doxycycline (2 µg/mL) was added on the first day of half medium change to induce TetO gene expression. The following day, puromycin (1 µg/mL) was added to start cell selection. Two days after the puromycin selection, a total of 5,000 HAs (ScienCell Research Laboratories, CA, USA) were added per device. After 24 hours, Ara-C (2 µM) was added to stop their proliferation. Half of the medium was changed twice a week with complete BrainPhys medium + 2 µg/mL doxycycline for 14 days. After that, half medium change was performed only with BrainPhys medium.

Five microfluidic devices were employed for each experimental condition (BIN1 KO vs WT both for spontaneous neuronal differentiation and Ascl1 induction) and two independent cultures were performed. To assess the time-course effect, neuron cultures were stopped at 4 and 6 weeks.

### Viral Transductions

Lentiviral constructs were produced by the Vect’UB platform within the TBM Core unit at University of Bordeaux, Bordeaux, France (CNRS UMS 3427, INSERM US 005). The lentiviral constructs used were TTA (Vect’UB; ID # 571) and TetO-Ascl1-Puro (Addgene, # 97329). Lentiviral infections were done in NPCs at P3 or P4. The viral constructs were transduced at a multiplicity of infection (MOI) of 2.5. In brief, NPCs were plated at a confluency of 1×106 cells per well of a 6-well plate. After 4 hours of plating the cells, appropriate volumes of each lentiviral construct were mixed in complete Neural Progenitor medium and 50 µl of the viral medium mix was then added to each well. We transduced the TTA construct at first in the hiNPCs. Following one passage, the TTA-transduced cells were transduced with the construct for ASCL1. Cells having both viral constructs were then further expanded for 1 or 2 passages before being used for differentiation into hiNs.

The iGluSnFR construct was an adeno-associated viral vector (BS11-COG-AAV8) sourced from Vigene Biosciences, MD, USA. The viral construct was transduced at a MOI of 5,000 at around 10 days of differentiation for the ASCL1-hiNs. Differentiation was allowed to continue for a duration of 4-6 weeks prior to MEA electrophysiology or imaging.

### Immunocytochemistry and Immunohistochemistry

Bidimensional (2D) cultures: All cells were fixed in 4% (w/v) paraformaldehyde (Electron Microscopy Sciences, Catalog # 15712) for 10 minutes in the imaging plates. Following, fixation, cells were washed thrice with PBS 0.1 M. Blocking solution (5% normal donkey serum + 0.1% Triton X-100 in PBS 0.1 M) was added to fixed cells at room temperature for 1 hour under shaking conditions. After the blocking step, primary antibodies were added to cells in the blocking solution and incubated overnight at 4°C. The following day, cells were washed with PBS 0.1 M thrice for 10 mins. Each. Alexa Fluor®--conjugated secondary antibodies in blocking solution were then incubated with the cells for 2 hours at room temperature under shaking conditions ensuring protection from light. Subsequently, 3 washes with 0.1 M PBS were done for 10 min each at room temperature under shaking conditions with protection from light. Hoechst 33258 solution was added during the second PBS wash. Cells were mounted with Aqua-Poly/Mount (Polysciences, Inc.) and imaged directly in the cell imaging plates. All images were acquired using an LSM 880 Confocal Scanning Microscope housed at the Imaging Platform of the Pasteur Institute, Lille. Duolink® Proximity Ligation Assays (PLA) was used to detect endogenous Protein-Protein Interactions. The following pairs of antibodies were used: anti-BIN1 (rabbit, 182562, abcam) and anti-Cav1.2 (mouse, 84814, abcam); or anti-BIN1 and anti-Cav1.3 (mouse, 85491, mouse). Other antibodies used for immunocytochemistry were: MAP2 (188006 and 188004, Synaptic Systems), Beta III Tubulin (MAB1637, Sigma-Aldrich), GFAP (AB5804, Millipore; and 173006, Synaptic Systems). All Alexa Fluor®-tagged secondary antibodies were sourced from Jacskon ImmunoResearch Europe Ltd.

Microfluidic Devices: Cultured induced neurons were fixed in 4% paraformaldehyde in PBS for 15 min at room temperature, washed three times with PBS, and permeabilized with 0.3% Triton X-100 in PBS for 5 min at room temperature. Cells were blocked in PBS containing 5% normal donkey serum for 1 h at room temperature before overnight incubation at 4°C with the following primary antibodies: MAP2 (188006, Synaptic Systems); HOMER1 (160004, Synaptic Systems), Synaptophysin (101011, Synaptic Systems), and GFAP (AB5804, Millipore). Cells were washed twice with PBS and incubated with the following secondary antibodies for 2h at room temperature: DyLight™ 405 Donkey Anti-Chicken (703-475-155, Jackson ImmunoResearch), Alexa Fluor 594 Donkey Anti-Guinea Pig (706-585-148, Jackson ImmunoResearch), Alexa Fluor 488 Donkey Anti-Mouse (715-545-151, Jackson ImmunoResearch) and Alexa Fluor 647 Donkey Anti-Rabbit (711-605-152, Jackson ImmunoResearch). Cells were rinsed three times with PBS and microfluidic devices were mounted with 90% glycerol.

Samples were imaged with a LSM 880 confocal microscope with a 63X 1.4 NA objective. Images were acquired at zoom 2 in z-stacks of 0.5 µm interval. Typically, 6 images were acquired per device from the synapse chamber near the postsynaptic chamber such the image contains multiple dendrites. Images were deconvoluted using the Huygens software (Scientific Volume Imaging, Netherlands).

Cerebral Organoids: Cerebral organoids were fixed in 4% PFA (w/v) for 30 min at 4°C followed by three washes with PBS 0.1 M. Cerebral organoids were then placed in sucrose solution (30% w/v) overnight before being embedded in O.C.T (Tissue-Tek). Embedded tissue was sectioned at 20 μm using a Cryostar NX70 Cryostat (Thermo Scientific) and mounted slides were stored at −80°C until immunostaining was performed. For immunostaining, tissue sections were brought to room temperature and then rehydrated with 3 washes with 0.1 M PBS, each for 5 mins. Slides were then washed once with PBS with 0.2% Triton X-100 for 15 mins. Tissue was blocked using 10% of donkey serum in PBS 0.1 M for 1 h at room temperature. After blocking, primary antibodies were added to 0.2 % Triton X-100 and 10% of donkey serum in PBS 0.1 M at appropriate dilutions and incubated overnight at 4°C. The next day, slides were washed with PBS 0.1 M 3 times for 5 min each with gentle shaking. Subsequently, slides were incubated with Alexa Fluor®-conjugated secondary antibodies in 0.2 % Triton X-100 and 10% of donkey serum in PBS 0.1 M for 2 h at room temperature in the dark. After secondary antibody incubation, slides were washed 3 times with PBS for 5 min with gentle shanking. Nuclei were visualized by incubating the tissue for 5 min with Hoechst 33258 stain in PBS 0.1 M. Sections were mounted using aqueous mounting medium (Polysciences). Images were acquired using an LSM 880 Confocal Scanning Microscope in concert with the ZEISS ZEN imaging software housed at the Imaging Platform of the Pasteur Institute, Lille. Image acquisition was done at 40X for the various cellular markers in Fig. 1. The antibodies used were MAP2 (188006, Synaptic Systems) and GFAP (AB5804, Sigma-Aldrich).

### Quantification of Synaptic Connectivity

Synaptic connectivity was quantified as previously described (Kilinc et al, 2020). Briefly, images were analyzed with Imaris software (Bitplane, Zürich, Switzerland) by reconstructing Synaptophysin I and HOMER1 puncta in 3D. The volume and position information of all puncta were processed using a custom Matlab (MathWorks, Natick, MA) program. This program assigns each postsynaptic spot to the nearest presynaptic spot (within a distance threshold of 11µm) and calculates the number of such assignments for all presynaptic puncta.

### Immunoblotting

Samples from the 2D cultures or brain organoids were collected in RIPA buffer containing protease and phosphatase inhibitors (Complete mini, Roche Applied Science) and sonicated several times at 60%-70% for 10 seconds prior to use for the immunoblotting analyses. Protein quantification was performed using the BCA protein assay (ThermoFisher Scientific). 10 μg of protein from extracts were separated in NuPAGE 4-12% Bis-Tris Gel 1.0mm (NP0321BOX, Thermo Scientific) or 3-8% Tri-Acetate gel (EA03755BOX, Thermo Scientific) and transferred on to nitrocellulose membranes 0.2μm (#1704158, Bio-Rad). Next, membranes were incubated in milk (5% in Tris-buffered saline with 0.1% Tween-20 (TBST)) or SuperBlock (37536, ThermoFisher Scientific) to block non-specific binding sites for 1 hour at room temperature, followed by several washes with TBST 0.1% or TNT 1x as washing buffers. Immunoblottings were carried out with primary antibodies overnight at 4°C under shaking condition. The membranes were washed three times in the washing buffer, followed by incubation with HRP-conjugated secondary antibodies for 2 hours at room temperature under shaking condition. The membranes were washed three times in washing buffer, and the immune reactivity was revealed using the ECL chemiluminescence system (SuperSignal, ThermoScientific) and imaged using the Amersham Imager 600 (GE Life Sciences). Optical densities of bands were quantified using the Gel Analyzer plugin in Fiji– ImageJ. The primary antibodies used for the immunoblots were as follows: BIN1 (ab182562,Abcam), APP C-terminal (A8717, Sigma-Aldrich), Tau (A002401-2, Agilent) Phospho-Tau(Clone: AT8) (MN1020,ThermoFisher Scientific), CaV1.3 (CACNA1D) (ACC-005, Alomone), CaV2.1 (CACNA1A) (ACC-001, Alomone), CaV2.2 (CACNA1B) (ACC-002, Alomone), CaV2.3 (CACNA1E) (ACC-006, Alomone), CaV1.2 (CACNA1C) (AGP-001 and ACC-003, Alomone), blocking peptide for Anti-CaV1.2 (CACNA1C) (BLP-CC003, Alomone) and β-ACTIN (A1978, Sigma-Aldrich). Secondary antibodies used for the immunoblots were Mouse-HRP (115-035-003, Jackson ImmunoResearch), Rabbit-HRP (111-035-003, Jackson ImmunoResearch), and Guinea pig-HRP (106-035-003, Jackson ImmunoResearch).

### Activity-dependent endosytosis assay

ASCL1-hiNs (n=9 cultures from each genotype) were subjected to 30 min of depolarization with 65 mM KCl or a mock treatment. Cells were then collected and pulled for endosomal fraction purification using the Minute™ Endosome Isolation and Cell Fractionation Kit (Invent Biotechnologies). Western blot was performed as described above.

### AlphaLISA measurements

Cell culture media samples for AlphaLISA measurements were collected at the endof the 3rd and 4th weeks of differentiation of the ASCL1-hiNs. Alpha-LISA kits specific for human Aβ1–X (AL288C, PerkinElmer) and Aβ1–42 (AL276C, PerkinElmer) were used to measure the amount of Aβ1–X and Aβ1–42 respectively in culture media. The human Aβ analyte standard was diluted in the BrainPhys medium. For the assay, 2 µL of cell culture medium or standard solution was added to an Optiplate-384 microplate (PerkinElmer). 21µL of 10X mixture including acceptor beads and biotinylated antibody was then added to the wells with culture media or standard solution. Following incubation at room temperature for an hour, 161µL of 1.25X donor beads was added to respective wells and incubated at room temperature for 1 hour. Luminescence was measured using an EnVision-Alpha Reader (PerkinElmer) at 680-nm excitation and 615-nm emission wavelengths.

### Calcium and iGluSnFR Imaging

Calcium imaging was performed in 2D cultures after 4 weeks (Ascl1-induced). Prior to imaging, the cells were incubated with Oregon Green™ 488 BAPTA-1 (OGB-1) acetoxymethyl (AM) (ThermoFisher Scientific) for 1 h. A 2.5 mM stock solution of the calcium-indicator dye was prepared in Pluronic™ F-127 (20% solution in DMSO) (ThermoFisher Scientific). 1 µL of the dye solution was added to 400 µL of fresh BrainPhys medium in each well of a 24-well cell imaging plate. Existing BrainPhys media from the wells of the plate was removed and kept aside while the calcium-indicator dye was incubated in fresh BrainPhys medium. After 1 h of incubation, the medium which was kept aside was replaced to each well. The 2D cultures were then ready to be imaged using a Spinning Disk Microscope (Nikon) housed at the Institut Pasteur de Lille, Lille, France using the MetaMorph imaging software.

For imaging the calcium activity, 1000 images were taken using a 20× long-distance objective, with 10 ms exposure time and 200 ms intervals. For each well, 5 random fields were chosen, and the cellular activity was recorded.

Cells transduced with iGluSnFR were directly imaged after 4 or 6 weeks of differentiation and 500 images were taken with the same imaging parameters. Up to 8 fields per well were filmed, each field containing at least one fluorescent transduced cell along with its processes.

### Analyses of Calcium Transients

All live recordings of neuronal calcium transients were first converted into .avi format after background subtraction using the FIJI software. Following the conversion, the videos were subsequently opened using the free software for data analyses of calcium imaging, CALciumIMagingAnalyzer (CALIMA) made available online by Fer Radstake (Eindhoven University of Technology, The Netherlands). Each video recording of a field of cells was first downscaled to 2X in terms of size with a 10X zoom and was checked for the frame average mode. Moreover, in this first detection stage, pre-set filter parameters were adjusted and applied to enable the detection of the maximum number of fluorescent cells in each field. In the analysis tab, detection of the average activity was checked and for pre-processing, a median of 3 was applied. All cells within the pre-set filter parameters are detected as regions of interest (ROIs) in the detection stage. Cell activity from all detected ROIs is then recorded. However, in the subsequent analysis stage, only cells showing spiking frequencies with a standard deviation of at least 2 or more were taken into consideration. Data in the form of detection spikes and the correlation (peak) are extracted and exported as .csv files.

### Electrophysiological recordings in 2D cultures and cerebral organoids

ASCL1-hiNs were cultured in microfluidic devices bound to multi-electrode arrays (256MEA100/30iR-ITO, Multi-Channel Systems, Germany). Extracellular action potentials were recorded in 5 different cultures for both genotypes at 2, 3, 4 and 6 weeks of differentiation using the MEA2100-256-System (Multi-Channel Systems). Before recordings, MEAs were let stabilize for 5 min on the headstage to reduce artifacts due to medium movement. Signals were recorded for 1 min, at 40 kHz sampling rate, using Multi Channel Experimenter 2.16.0 software (Multi-Channel Systems). Electrical activity in cerebral organoids was recorded using 256-6wellMEA200/30iR-ITO (Multi-Channel Systems, Germany). Briefly, 5-6-month-old cerebral organoids were mounted onto MEAs and kept for 2 h in complete Brainphys medium. Then, MEAs were placed on the headstage and let stabilize for 5 min before recordings. Signals were recorded for 5 min, at 10 kHz sampling rate using Multi-Channel Experimenter 2.16.0. For rescue experiments using a calcium channel blocker, ASCL1-hiNs were cultured on MEA 96-well plates (CytoView MEA 96, Axion Biosystems, USA). Extracellular action potentials were recorded in 3 independent cultures for either genotype in the presence of 50nM nifedipine (Tocris Bioscience) or vehicle using the MaestroPro (Axion Biosystems, Inc, USA). Before recordings, MEAs were let stabilize for 5 min on the MaestroPro at 37°C and 5% CO_2_. Signals were recorded for 3 min, at 12.5 kHz sampling rate, using AxIS Navigator software (Axion Biosystems).

Spikes were detected using a fixed amplitude threshold of 5.5 and 4.5 standard deviations (for the 2D and 3D cultures, respectively) of the high-pass filtered (>300 Hz) signal for positive-and negative-going signals. The detection included a dead time of 3 ms to account for the refractory period of action potentials. Quantification of the number of detected spikes (MUAs) and spike bursts (defined as at least 5 spikes within 50 ms) was performed using Multi-Channel Analyzer 2.16.0 software (Multi-Channel Systems).

### Spike sorting and temporal structure of spontaneous activity

Channels containing detected waveforms were manually processed offline for spike waveform separation and classification using Offline Sorter v3 (Plexon, USA). Briefly, we applied principal component analysis (PCA) to cluster spike waveforms of similar morphologies. Using this approach, we identified from 2 to 10 well-isolated units per channel, and therefore, we considered this single-unit activity (SUA). For each SUA, we computed the average firing rate, the signal-to-noise ratio, the peak-to-trough amplitude and duration, the average power (square amplitude of the average waveform), the mode of the interspike interval distribution, and their firing patterns. It has been demonstrated that dissociated neuronal cultures can develop complex discharge structures [47]. Here, we considered burst activity if the SUA presents periods of high-frequency discharges interspersed by regular or no discharges at all. Operationally, a burst must have at least 3 spikes within 100 ms and 200 ms intervals, for the interval between the first and the second, and the second and the third discharge, respectively. After the third spike, the maximal interval to consider a discharge part of the burst was 200 ms. Thus, we computed the SUA that presented bursts, the number of bursts (i.e., the burst frequency), the average burst duration, the number of spikes within each burst, the average burst frequency, and the inter-burst interval. To evaluate the temporal structures of spike trains, we computed the array-wide spike detection rate (ASDR), which is the number of spikes detected per unit of time, summed over all electrodes in the array. This method is commonly used in the literature to demonstrate synchronous activity (aka, bursts) in MUA data (Wagenaar, 2006).

### snRNA-seq Library Preparation

Nuclei isolation and Hash-tagging with oligonucleotides steps were realized on ice with pre-cold buffers and centrifugations at 4°C. 6.5-month-old BIN1 WT, HET, and KO organoids were processed as previously (Lambert et al., 2022). 4-week-old cultured ASCL1-induced BIN1 WT and KO 2D cultures were washed in the imaging plate wells with 500 µL of Deionized Phosphate Buffer Saline 1X (DPBS, GIBCO™, Fisher Scientific 11590476). Cells were resuspended with wide bore tips in 500 μL Lysis Buffer (Tris-HCL 10mM, NaCl 10mM, MgCl2 3mM, Tween-20 0,1%, Nonidet P40 Substitute 0,1%, Digitonin 0,01%, BSA 1%, Invitrogen™ RNAseout™ recombinant ribonuclease inhibitor 0,04 U/μL). Multiple mechanical resuspensions in this buffer were performed for a total lysis time of 15 mins., 500 μL of washing buffer was added (Tris-HCL 10mM, NaCl 10 mM, MgCl2 3 mM, Tween-20 0.1%, BSA 1%, Invitrogen™ RNAseout™ recombinant ribonuclease inhibitor 0,04 U/μL) and the lysis suspension was centrifuged 8 mins. at 500 g (used for all following centrifugation steps). Nuclei pellets were washed tree times with one filtration step by MACS pre-separation filter 20μm (Miltenyi Biotec). Nuclei pellets were resuspended in 100 μL of staining buffer (DPBS BSA 2%, Tween-20 0.01%), 10 μL of Fc blocking reagent HumanTruStainFc™ (422302, Biolegend) and incubated 5 min at 4°C. 1μl of antibody was added (Total-Seq™-A0453 anti-Vertebrate Nuclear Hashtag 3 MAb414 for the WT and Total-Seq™-A0454 anti-Vertebrate Nuclear Hashtag 4 MAb414 for the KO, 97286 and 97287 respectively, Biolegend) and incubated 15 mins. at 4°C. Nuclei pellets were washed three times in staining buffer with one filtration step by MACS pre-separation filter 20 μm (Miltenyi Biotec) to a final resuspension in 300 μL of staining buffer for Malassez cell counting with Trypan blue counterstaining (Trypan Blue solution, 11538886, Fisherscientific). Isolated nuclei were loaded on a Chromium 10X genomics controller following the manufacturer protocol using the chromium single-cell v3 chemistry and single indexing and the adapted protocol by Biolegend for the HTO library preparation. The resulting libraries were pooled at equimolar proportions with a 9 for 1 ratio for Gene expression library and HTO library respectively. Finally, the pool was sequenced using 100pb paired-end reads on NOVAseq 6000 system following the manufacturer recommendations (Illumina).

### snRNA-seq Dataset Preprocessing

Unique Molecular Index (UMI) Count Matrices for gene expression and for Hash Tag Oligonucleotide (HTO) libraries were generated using the CellRanger count (Feature Barcode) pipeline. Reads were aligned on the GRCh38-3.0.0 transcriptome reference (10x Genomics). Filtering for low quality cells according to the number of RNA, genes detected, and percentage of mitochondrial RNA was performed. For HTO sample, the HTO matrix was normalized using centered log-ratio (CLR) transformation and cells were assigned back to their sample of origin using HTODemux function of the Seurat R Package (v4) [48]. Then, normalizations of the gene expression matrix for cellular sequencing depth, mitochondrial percentage and cell cycle phases using the variance stabilizing transformation (vst) based Seurat:SCTransform function were performed.

### snRNA-seq datasets integration and annotation

To integrate the datasets from independent experiments, the harmony R package (https://github.com/immunogenomics/harmony) was used. In order to integrate the datasets, the SCTransform normalized matrices was merged and PCA was performed using Seurat::RunPCA default parameter. The 50 principal components (dimensions) of the PCA were corrected for batch effect using harmony::RunHarmony function. Then, the 30 first batch corrected dimensions were used as input for graph-based cell clustering and visualization tool. Seurat::FindNeighbors using default parameters and Seurat::FindClusters function using the Louvain algorithm were used to cluster cells according to their batch corrected transcriptomes similarities. To visualize the cells similarities in a 2-dimension space, the Seurat::RunUMAP function using default parameter was used. Cell clusters were then annotated based on cell type specific gene expression markers.

### Differential gene expression and GO enrichment analyses

For snRNA-seq experiment reproduce 2 times or less (i.e. organoid and nifedipine experiments), gene expression within each main cell type was compared between conditions of interest using Wilcoxon test on the SCTransform normalized gene expression matrix. For snRNA-seq experiment reproduce more than 2 times (BIN1 KO vs WT hiNs comparison), Pseudobulk differential expression analysis was performed using the package DESeq2 (CRAN). For each main cell type, the cell level gene expression matrix was first aggregate at the sample level using Matrix:: aggregate.Matrix function. Genes detected in more than 100 cells or in 10% of the cells was test for differential expression using DESeq function, including in the model the experiment batch as covariate.

GO enrichment analysis on the differentially expressed genes was performed using the gost function of the gprofiler2 R package (CRAN), using default parameter, or, if specified using fast gene-set enrichement analysis (FGSEA, [49]) package (Bioconductor).

### Activity-related genes (ARGs) signature enrichment analysis at single cell resolution

To study enrichment for activity-related genes (ARGs) signature across cerebral organoid cells, the CellID R package (https://github.com/RausellLab/CelliD) was used. ARGs obtained from Tyssowski et al. (2018) and Hravtin et al. (2018) (supplementary Table 7), were translated to the corresponding human gene name with the help of the biomaRt package using the respective Ensembl references. Then, the CellID::RunMCA was used to extract cell-specific gene signature and hypergeometric test was performed to test enrichment for ARGs in these cell signatures. To test the differential proportion of ARGs enriched cells in BIN1 deleted organoid compared to WT organoid, chi-squared test was performed.

### Comparative analysis with specific DEGs in AD brains

To compare the transcriptomic change observed in BIN1 deleted cerebral organoid with those observed in AD brain we used the dataset available for download at Synapse.org under the Synapse ID syn21788402 [35]. The raw gene expression matrix was normalized using Seurat::SCTransform and differential expression analysis was performed within each neuronal cell type using Wilcoxon test as used for our organoid dataset. AD related DEGs, thus, obtained were compared with our BIN1 related organoid DEGs in every cell type. To this end, the enrichment for AD-related DEGs in BIN1-related DEGs was tested using hypergeometric test. The background for this test was defined as all genes detected in both datasets. The p-value of this test was used as metrics to compare the significance of the gene overlap between neuronal cell types.

### Statistical analyses

Statistical analyses were performed using GraphPad Prism version 8.0.0 (GraphPad Software, San Diego, California USA, www.graphpad.com) and R 4.2.0 (R Core Team, 2022, https://cran.r-project.org/bin/windows/base/old/4.2.0/). Bar plots show mean ± SD and individual values. Box plots show 1-99 percentile. Statistical tests and p values are indicated in figure legends.

**Sup. Figure 1:**
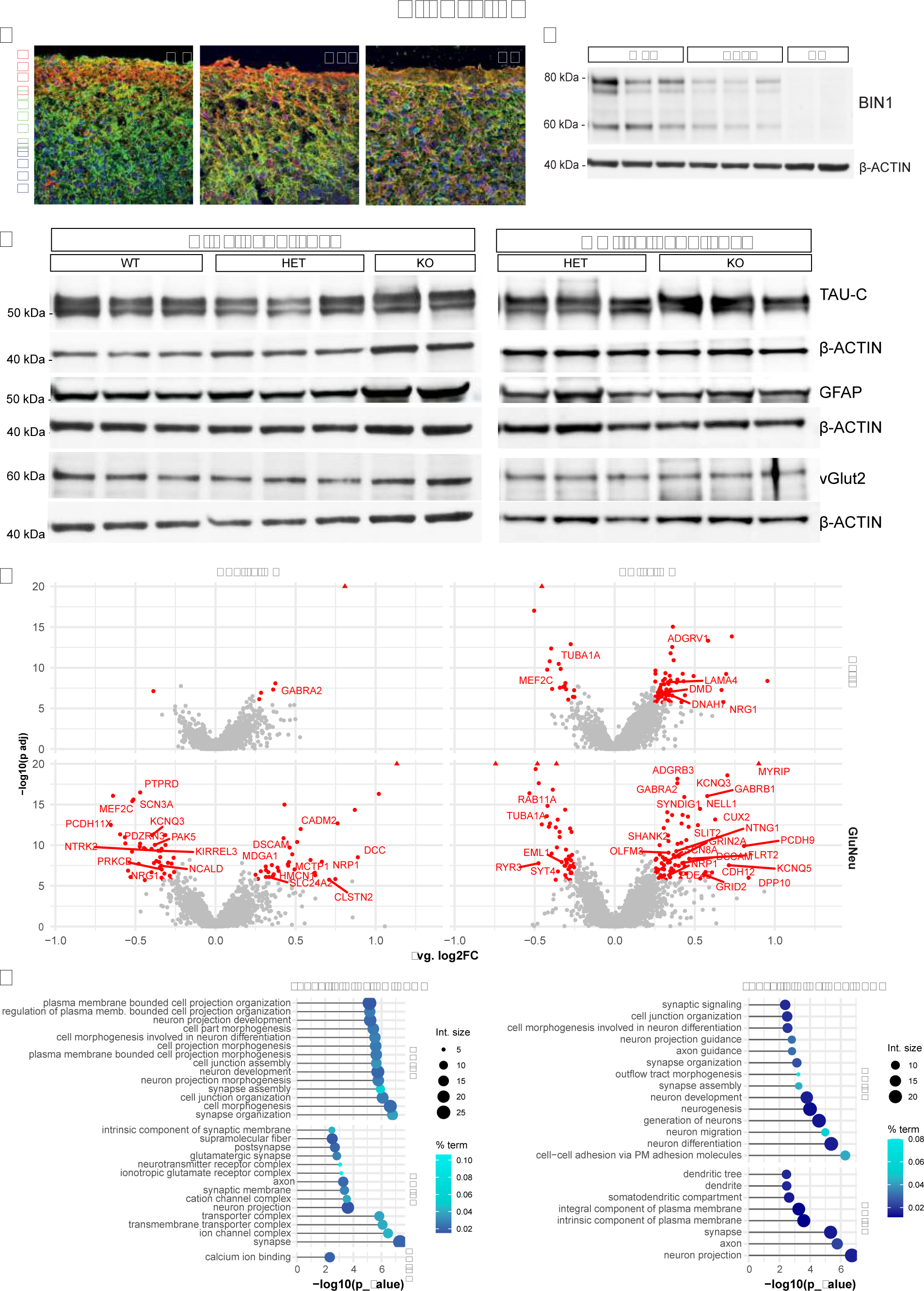
Cellular and molecular characterization of 6.5-month-old cerebral organoids. (A) Immunohistochemistry for GFAP (red), MAP2 (green) and DAPI (blue) in 6.5-month-old *BIN1* WT, HET and KO COs. (B) Western blots showing the isoforms of BIN1 detected in WT, WT, the reduced expression in HET, and the absence of BIN1 protein in KO COs. (C) Western blots showing the expression of neuronal (TAU and vGLUT2) and astrocytic (GFAP) proteins in 14 individual COs at 190 days, including some of the BIN1 WT, HET and KO samples used for snRNA-seq. Note the low variability in protein expression. (D) Volcano plots representing DEGs comparing HET vs WT or KO vs WT in astrocytes and glutamatergic neurons. DEGs with adjusted p-value <0.05 and |log2FC| >0.25 are shown in red. Gene labels are shown for top 10 genes in terms of log2FoldChange and p-value. (E) (H) Functional enrichment analysis showing the top 15 significantly enriched GOs for DEGs identified in the *BIN1* KO vs WT or HET vs WT glutamatergic neurons. GO: gene ontology; BP: biological processes; CC: cellular components; MF: molecular function.

**Sup. Figure 2:**
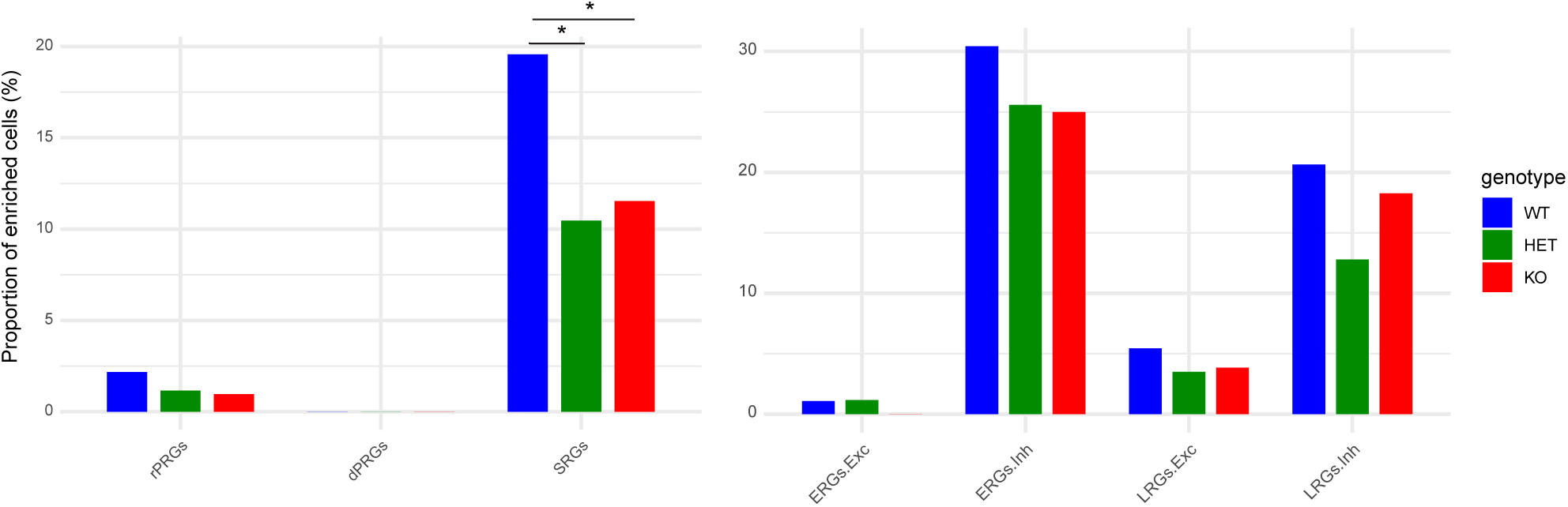
Reduced BIN1 expression does not enhance activity-dependent transcription in GABAergic neurons. A-B) Proportions of GABAergic neurons enriched for the different ARG signatures according to genotype (*p<0.05; **p<0.01; ***p<0.001; Chi-squared test).

**Sup. Figure 3:**
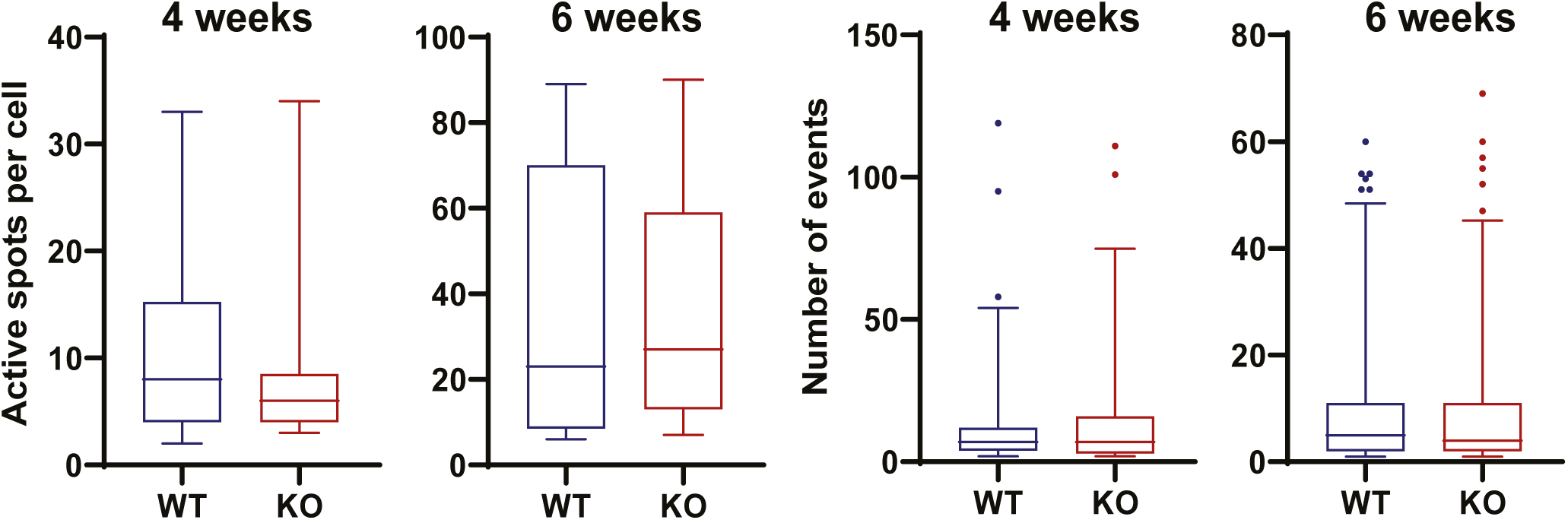
Normal glutamatergic transmission in BIN1 KO ASCL1-hiNs. **Box plots** show the number of active spots per neuron and number of events detected by time-lapse video-microscopy in 4- or 6-week-old ASCL1-hiNs cultures transduced with the glutamate sensor iGLUSnFr (4 weeks: n= 378 BIN1 WT and 266 BIN1 KO ASCL1-hiNs; 6 weeks: n= 685 BIN1 WT and 629 BIN1 KO ASCL1-hiNs).

**Sup. Figure 4:**
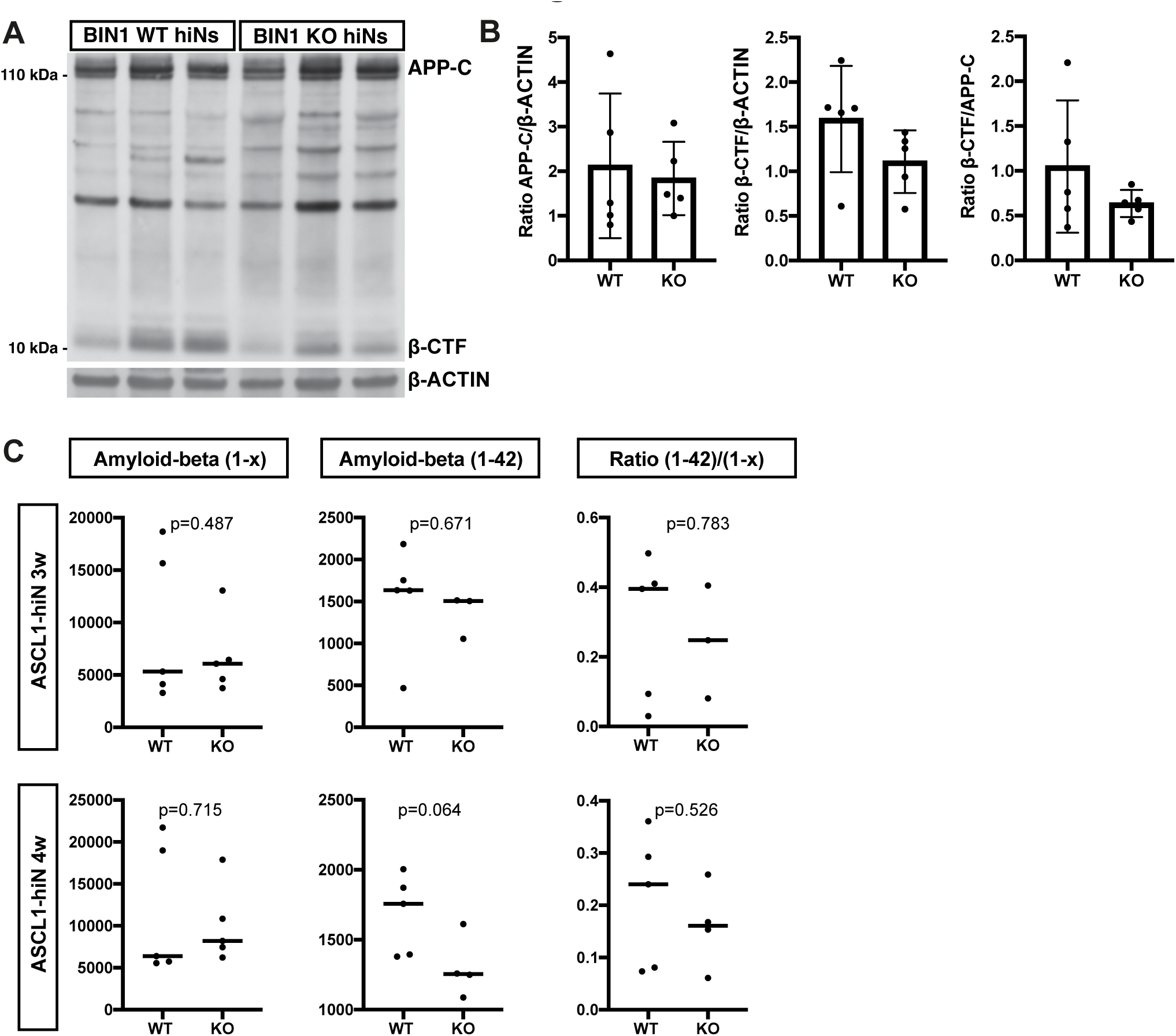
Normal APP processing in BIN1 KO ASCL1-hiNs. (A) Western blots showing the expression of APP full-length, β-CTF and β-ACTIN at 4 weeks. (B) Quantification of the ratios APP/β-ACTIN, β-CTF /β-ACTIN and β-CTF/APP (n = 5 for each genotype). (C) Quantification of soluble Aβ_1-x_, Aβ_1-42_ and the ratio Aβ_1-42/_ Aβ_1-x_ in ASCL1-hiNs cultures at 3 and 4 weeks.

**Sup. Figure 5:**
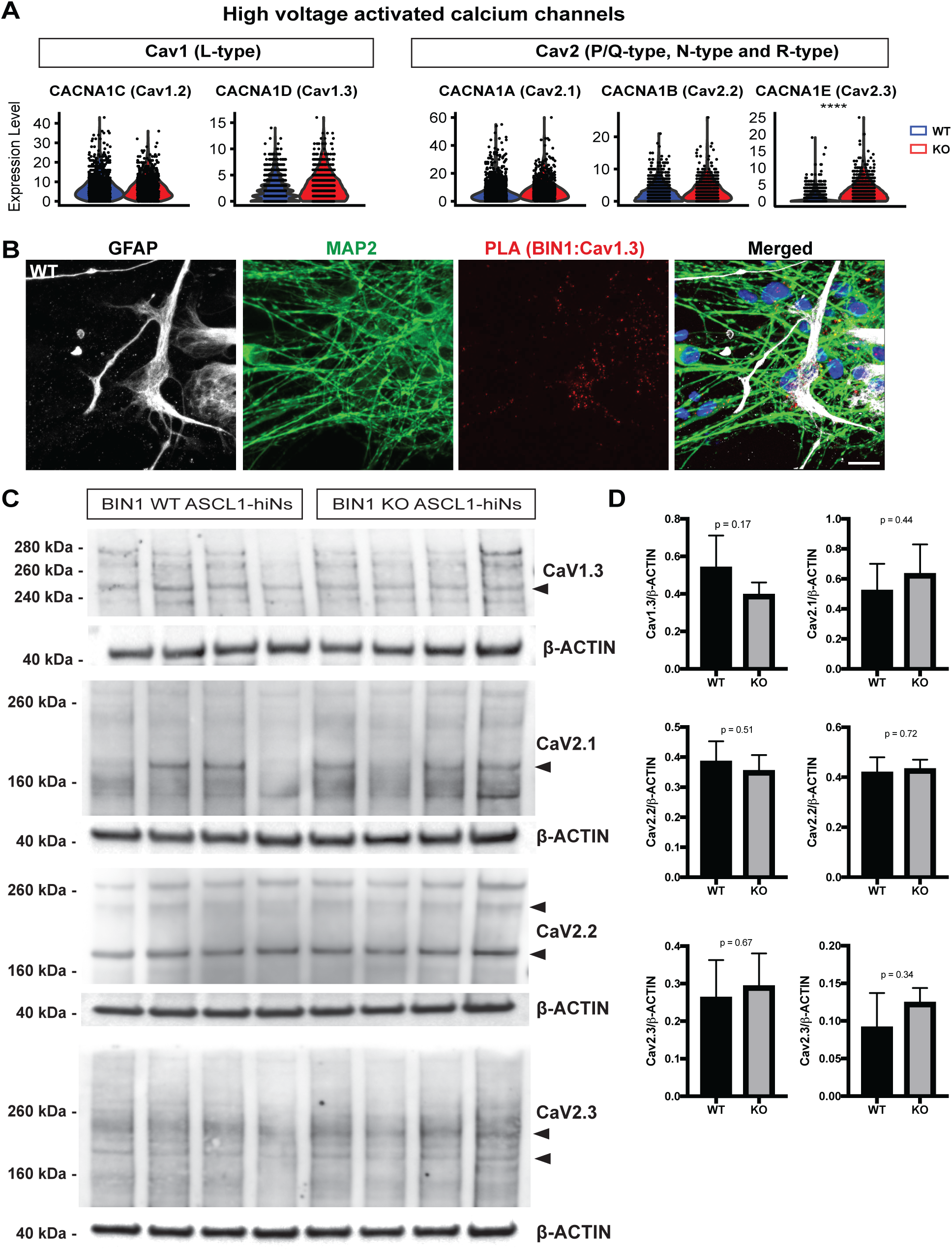
Expression of voltage-gated calcium channels in ASCL1-hiNs. (A) Violin plots showing the mRNA levels of Cav1 and Cav2 members of the voltage-gated calcium channel families L-type, P/Q-type, N-type and R-type detected in ASCL1 hiNs. (B) Images showing PLA spots using anti-BIN1 and anti-Cav1.3 antibodies in 4-week-old *BIN1* WT hiNs. Cells were also immunolabeled for the neuronal marker MAP2 (green), the astrocyte marker GFAP (white), and stained with DAPI (blue). (C) Western blots showing the expression of Cav1.3, Cav2.1, Cav2.2, Cav2.3 and β-ACTIN in 4-week-old ASCL1h hiNs. (D) Quantification of protein expression.

**Sup. Figure 6:**
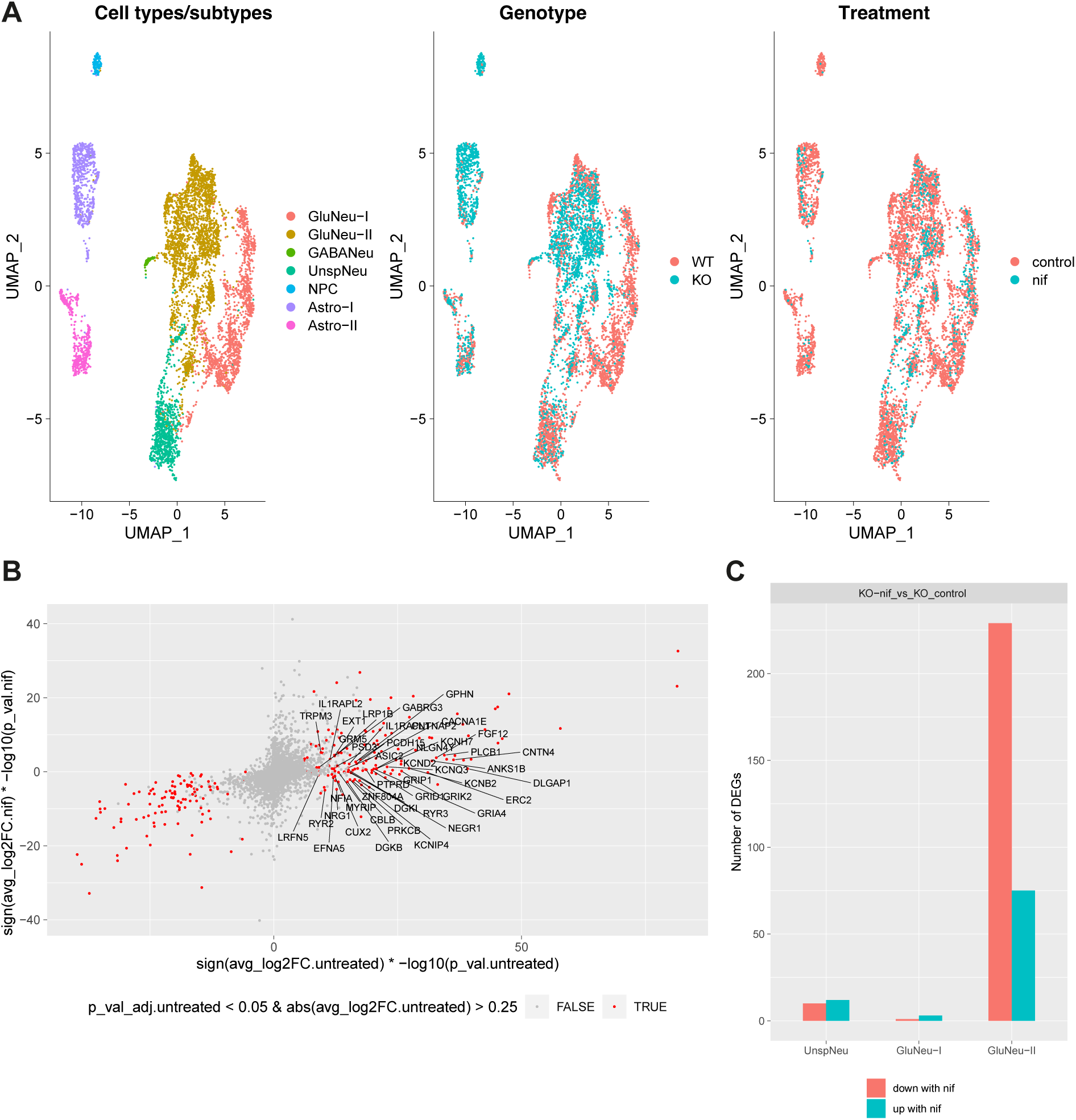
snRNAseq of nifedipine treated ASCL1-hiNs cultures. (A) UMAP of the n=2 nifedipine experiments integrated with the other hiNs snRNA-seq experiments. (B) correlation between the differential gene expression in untreated *BIN1* KO + vehicle vs WT (x axis, DEGs are plotted in red) and in *BIN1* KO + nifedipine (nif) vs WT. Calcium- and synapse-related genes upregulated in *BIN1* KO + vehicle vs WT are labeled if the genes are not upregulated in KO + nif vs WT. (C) Plot showing the total number of DEGs identified in *BIN1* KO ASCL1-hiNs treated or not with nifedipine.

